# Automated detection, morphometric measurement, and size-class determination of harbor seals (Phoca vitulina) from UAV orthomosaics: a cascaded deep-learning pipeline with stratified validation across platforms, sensors, altitudes, and substrates

**DOI:** 10.64898/2026.10.01.755886

**Authors:** Ivan A. Usatov, Vladimir N. Burkanov, Alexey V. Altukhov, A. Bowker, Chad Nordstrom, Strahan Tucker, Irina S. Trukhanova

**Affiliations:** North Pacific Wildlife Consulting LLC, Lake Forest Park, Washington, United States of America; Marine Mammal Laboratory, Alaska Fisheries Science Center, National Marine Fisheries Service, National Oceanic and Atmospheric Administration, Seattle, Washington, United States of America; Pacific Biological Station, Fisheries and Oceans Canada, Nanaimo, British Columbia, Canada

## Abstract

Reliable demographic data are critical for seal population assessment and management, yet remain challenging to obtain from drone surveys without automated tools. Using data from harbor seal (*Phoca vitulina*) haulout surveys conducted in British Columbia, Canada, in 2023–2024, we developed and tested an automated pipeline that integrates detection, morphometric extraction, and size-class classification of seals into a single system. The pipeline combines cascaded DeepLabV3+ segmentation (water removal, colony-level seal detection, and individual body-contour refinement), principal component analysis (PCA)-based extraction of body length and maximum width, and three-class size-class determination based on body length (pups, juveniles, subadults/adults). We evaluated the pipeline on 2,483 manually annotated seals across four drone platforms (DJI Mavic 3 Thermal, Mavic 2 Pro, Mini 3 Pro and Mini 2), five substrate types, and two sensor modalities (RGB, thermal infrared). Aggregate detection F1 was 0.914. Orthomosaic geometric accuracy, quantified as the RMSE of calibration-rod length on the orthomosaic, ranged from 1.0 cm to 18.0 cm depending on the platform–altitude–substrate combination, a spread that aggregate metrics would mask. Relative to manual annotation on the same orthomosaics, automated RGB body lengths showed near-zero bias (−0.6 cm; RMSE = 12.5 cm), whereas automated thermal lengths were systematically shorter (bias −19.9 cm, RMSE = 24.1 cm), establishing the unsuitability of thermal imagery for automated quantitative pinniped morphometry. For RGB imagery, threshold classification by body length reached an overall accuracy (OA) of 78.7%; for individuals outside the uncertainty zones (defined as ±2 × the geometric RMSE on calibration rods around each class boundary), OA reached 90.9%. Within the RGB subset, OA was 78.9% at 20 m altitude, declining to 75.0% at 40 m where geometric error dominated. The pipeline enables automated demographic monitoring of pinniped haulouts with quantitatively characterized uncertainty across survey conditions.

## 1. Introduction

Unoccupied aerial vehicles (UAVs) have transformed wildlife monitoring over the past decade, providing high-resolution imagery with reduced disturbance, lower operational costs, and greater deployment flexibility compared to crewed aircraft [1–3]. For pinnipeds, which aggregate at terrestrial haulouts in numbers that must be surveyed repeatedly and over large spatial extents, drones offer four key capabilities: repeat and on-demand deployment, high-resolution spatial coverage, morphometric photogrammetry, and compatibility with computer vision and deep learning applications [4]. These capabilities have driven rapid adoption of UAV platforms by agencies responsible for pinniped population assessment, yet the data-processing infrastructure required to realize their full potential has not kept pace with the rate of image acquisition.

Aerial surveys have historically been the primary method for estimating pinniped abundance at haulout sites [5]. While crewed-aircraft programs generate manageable volumes of imagery, a single UAV flight at survey altitude can produce hundreds to thousands of overlapping frames that must be stitched into georeferenced orthomosaics before any ecological information can be extracted. When monitoring programs require repeated coverage of dozens of sites across seasons and years - as is the case for species such as the harbor seal (*Phoca vitulina* Linnaeus, 1758), which is surveyed by Fisheries and Oceans Canada (DFO) at sites throughout British Columbia - the resulting data volumes quickly exceed the capacity of manual interpretation. Scaling from image acquisition to population-level inference therefore demands automated processing pipelines that can detect individuals, extract morphometric measurements, and assign demographic classes with quantified and reproducible error.

Deep learning methods have achieved high accuracy in automated pinniped detection from aerial imagery [6,7], and UAV-based photogrammetry has demonstrated potential for non-invasive morphometric assessment across several species, including the Australian sea lion (*Neophoca cinerea*) [8], the harbor seal [9], and the Pacific walrus (*Odobenus rosmarus divergens*), for which body mass of reproductive-age females can be estimated from two-dimensional imagery with errors comparable to those of three-dimensional methods [10]. UAV photogrammetry has also been applied to assess abundance and size composition of large spotted seal (*Phoca largha*) aggregations at coastal haulouts in the southwestern Bering Sea [11].

However, in all of these studies, morphometric parameter extraction was performed manually or semi-automatically, creating a persistent bottleneck between raw image capture and usable demographic data. This dependence on manual processing likely contributes to the fact that the vast majority of published automated detection studies remain limited to aggregate counts and lack the age-class resolution required for management applications that depend on demographic structure — such as distinguishing stable populations from those experiencing cryptic declines in recruitment or body condition [9,10,12]. Manual age determination from body length on aerial photographs has been attempted [13] but is extremely labor-intensive; our preceding work [11] required manual measurement of 4,057 individual spotted seals and, while confirming the value of UAV photogrammetry for size-composition analysis, pointed to the need for an automated pipeline with calibrated error assessment.

A critical gap therefore persists: while automated detection is common, no published study has yet demonstrated a fully automated pipeline that integrates detection, morphometric extraction, and size-class classification of pinnipeds from raw UAV orthomosaics—the demographic products monitoring agencies ultimately require. Additional tool-development challenges include: (1) insufficient characterization of measurement error stratified by platform–altitude–substrate interactions under real field conditions; (2) the lack of a quantitative link between measurement error and size-class misclassification, including the width of the uncertainty zones at class boundaries; (3) limited evaluation of processing performance on the heterogeneous rocky substrates typical of many haulout sites, where background complexity degrades both detection and segmentation; and (4) the absence of a systematic comparison between thermal infrared and RGB sensor modalities for quantitative morphometry within a single automated framework.

The Strait of Georgia, British Columbia, Canada, was selected as the study area because it offers an ideal testbed for pipeline development. The region supports the highest density of *P. vitulina* on the British Columbia coast at 10.5 seals/km, with a population of approximately 39,000 individuals (95% CI: 33,400–45,200) [14,15] that stabilized in the mid-1990s, providing a large and consistent source of training and validation imagery. Haulout sites encompass a variety of substrate types (rocky slopes, boulders, reefs, plateaus, sand beaches), permitting systematic evaluation of substrate effects on detection and measurement accuracy - a key requirement for tools intended to generalize across survey programs. The logistical accessibility of the haulouts and the availability of long-term aerial survey data from DFO (since 1966) provide the reference infrastructure necessary for tool validation and benchmarking.

Seasonal variation in haulout composition further supports pipeline evaluation across contrasting demographic scenarios. The pupping season of *P. v. richardii* in southern British Columbia extends from late May through July, with a peak in June; lactation lasts 4–6 weeks, after which pups transition to independent foraging. Adult molt occurs in August–September, when the proportion of animals hauled out is at its annual maximum [16,17]. September surveys therefore yield the highest counts and the densest aggregations. The two periods also differ in composition, although less straightforwardly than the count maximum suggests: June surveys encounter dependent pups alongside their mothers, whereas by September that cohort has weaned and hauls out independently, and haul-out probability itself varies with age and sex, so neither period samples the population’s size structure without bias [18,19].

Here we develop and validate an AI-based processing pipeline that integrates detection, two-dimensional morphometric measurement, and size-class classification of phocid seals, using *P. vitulina* as a model system, within a single automated workflow applied to UAV orthomosaic imagery. We systematically stratify measurement error and classification accuracy across multiple platforms, altitudes, substrates, and sensor modalities, with the aim of producing an operationally deployable tool that bridges the gap between drone image acquisition and the demographic data products required for effective pinniped monitoring and management.

## 2. Materials and Methods

### 2.1. Study area and field data collection

We conducted aerial surveys of *P. vitulina* at 20 haulout sites in British Columbia, Canada (Fig 1) between May 3 and July 20, 2023, and September 16–19, 2024. Four multirotor drone platforms were used: DJI Mini 2 (Mn2), DJI Mini 3 Pro (Mn3), DJI Mavic 2 Pro (M2P), and DJI Mavic 3 Thermal (M3T). All platforms carried built-in RGB cameras; the M3T was additionally operated in long-wave infrared (LWIR) mode. All platforms relied solely on their onboard single-frequency GNSS receivers, which track only the L1 civil band (1575.42 MHz); no ground control points (GCPs) or real-time kinematic (RTK) base station were used, consistent with the conditions of most routine UAV haulout surveys flown with consumer-grade drones.

**Fig 1.**
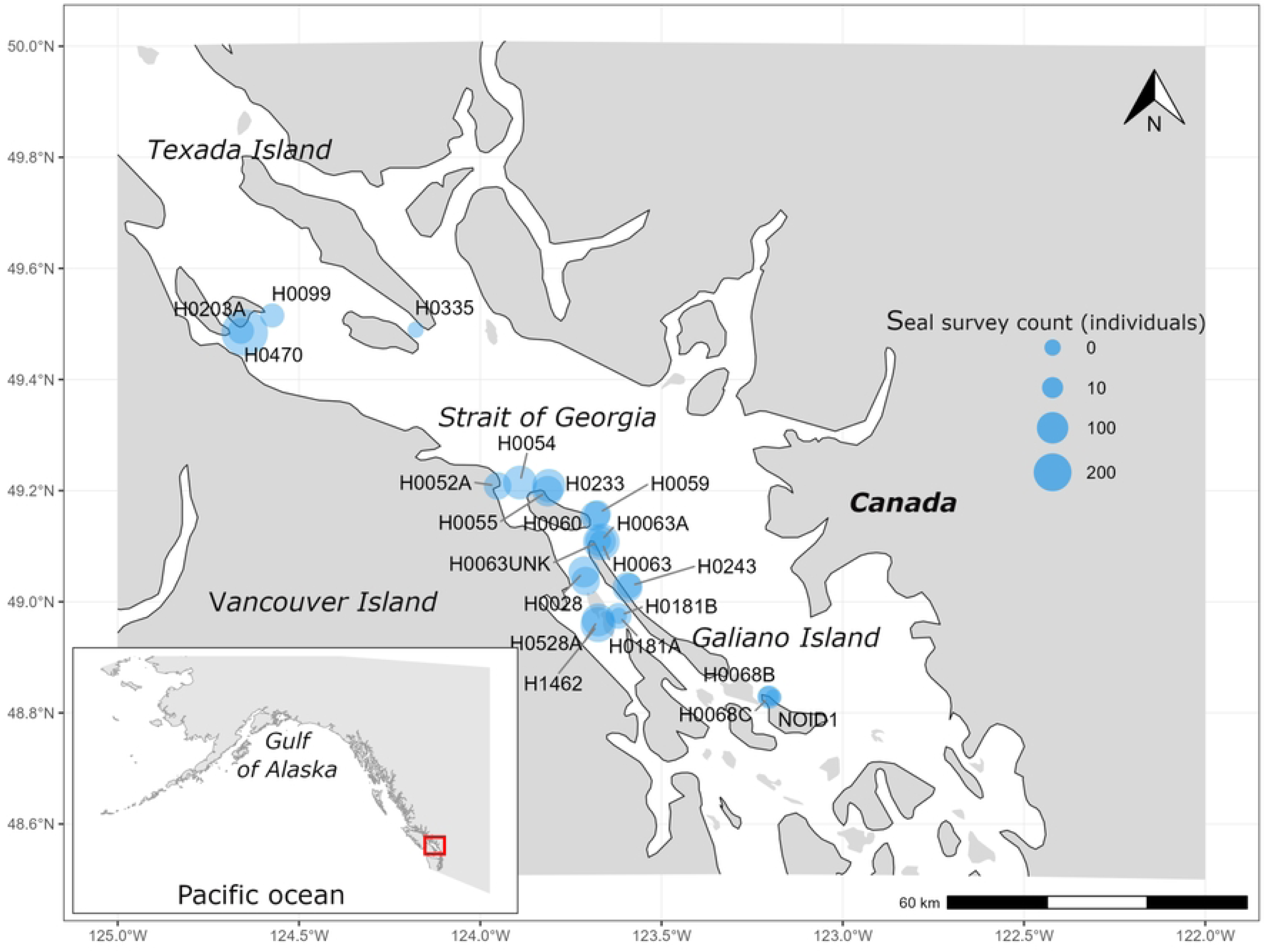
**Map of *P.*** *vitulina* haulout survey sites in the Strait of Georgia, British Columbia, Canada. Circle size is proportional to the number of detected individuals at each site. Insert shows the study area location on the Pacific coast of North America.

Flight altitude ranged from 20 to 40 m depending on colony size and behavioral response; the standard protocol was 20 m. Image overlap was at least 70% in both directions. Approximately 30,000 images were acquired, from which 53 unique orthomosaics were reconstructed: 31 seal-survey orthomosaics (animals measured) and 23 calibration orthomosaics bearing control rods (22 retained for geometric analysis and one excluded by the coverage-area criterion; S1 Table), with one orthomosaic (site H0054) carrying both seals and rods and therefore shared between the two sets. A further 10 harbor-seal orthomosaics from this study were used to train the segmentation models (Section 2.2). Wooden calibration rods of four standard lengths (0.46, 0.96, 1.21, and 1.36 m) were placed on the substrate surface as independent control markers for evaluating orthomosaic geometric accuracy (Fig 2). They were laid in the orientation permitted by the local micro-relief and were not levelled instrumentally. The rod lengths were selected to bracket the measurement range relevant to size-based classification of *P. vitulina*: the shortest rod (0.46 m) provides a sub-body-size reference for assessing small-object accuracy; the 0.96 m rod approximates the body length of weaned pups at ∼3 months of age (95–105 cm) [20,21]; the 1.21 m rod falls within the young-of-year/juvenile range (110–130 cm) [22,23]; and the 1.36 m rod corresponds to the juvenile/subadult boundary region (130 cm threshold used in this study; S2 Table).

**Fig 2.**
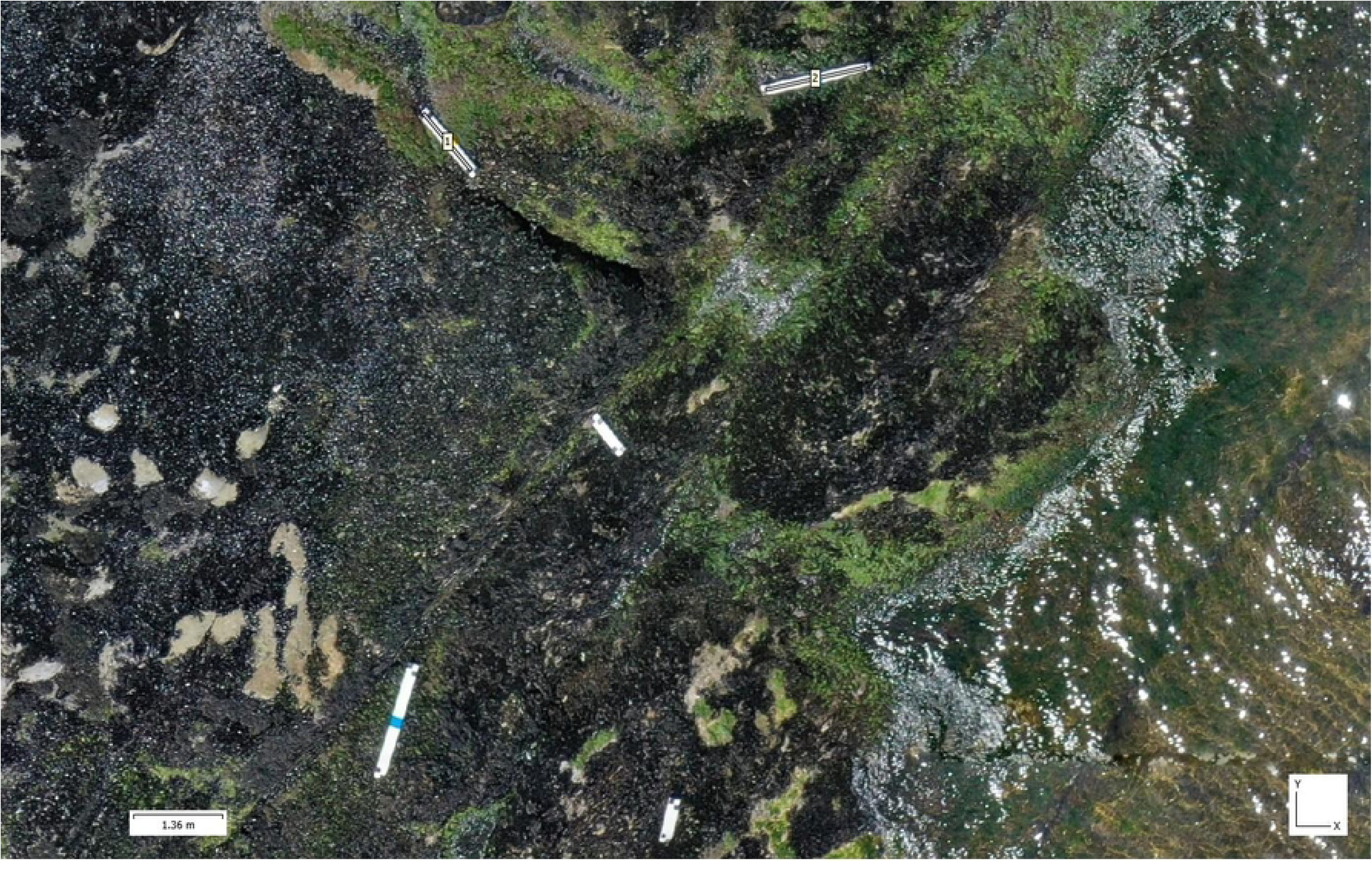
**Calibration rods of known length placed on rocky substrate before a drone survey for independent assessment of orthomosaic geometric accuracy.**

Orthomosaics were constructed in Agisoft Metashape (version 2.3.0) following a standard photogrammetric workflow: image alignment, dense point cloud generation, digital surface model creation, and orthomosaic production. Pixels classified as water by the water-removal model were masked before orthomosaic construction. This masking served two purposes. The principal one concerned morphometric reliability: a submerged seal, even when clearly visible from above, is at an unknown depth, so refraction at the air–water interface and the orthorectification error for an object lying below the reconstructed surface make its apparent length unreliable. Measurements were therefore restricted to animals fully hauled out on exposed substrate, and any individual still in the water, including one visible beneath the surface in a retained shallow area, was excluded from measurement. Masking removed open-water individuals directly and, secondarily, addressed a photogrammetric problem: the surf-zone water surface moves continuously between overlapping images and cannot be reconstructed reliably.

All linear measurements were converted from pixels to meters using the ground sample distance (GSD) derived from image metadata (flight altitude and camera parameters). Haulouts were associated with five substrate types: slopes, boulders, plateaus, reefs, and white sand beach.

All morphometric measurements were performed on SfM-derived orthomosaics rather than on individual aerial photographs [24–26].

### 2.2. Data annotation and calibration

Manual annotation was performed by trained observers using the custom-built software MaskCorrector (openly available at github.com/ivan-npwc/MaskCorrector) and the Agisoft Metashape environment. Each visible individual was delineated as a binary polygon mask coinciding with the outer body contour. Reference body-length and body-width measurements followed standard mammalogical protocols [27]. Body length was defined as the distance from the snout tip to the tail tip in dorsal projection, measured along the dorsal midline (defined as the longitudinal line of the dorsal silhouette lying equidistant from the left and right body outlines at each point along the animal, not the projection of the vertebral column), which follows the lateral curvature of the trunk. Width was the maximum transverse dimension measured perpendicular to the longitudinal axis (Fig 3). A formal inter-observer agreement assessment was not conducted. Published validations of manual measurements on drone orthomosaics for pinnipeds consistently report high reproducibility (ICC > 0.99) [8] and mean inter-observer discrepancy of 1–3 cm, which we consider sufficiently reliable for the purpose of this study.

**Fig 3.**
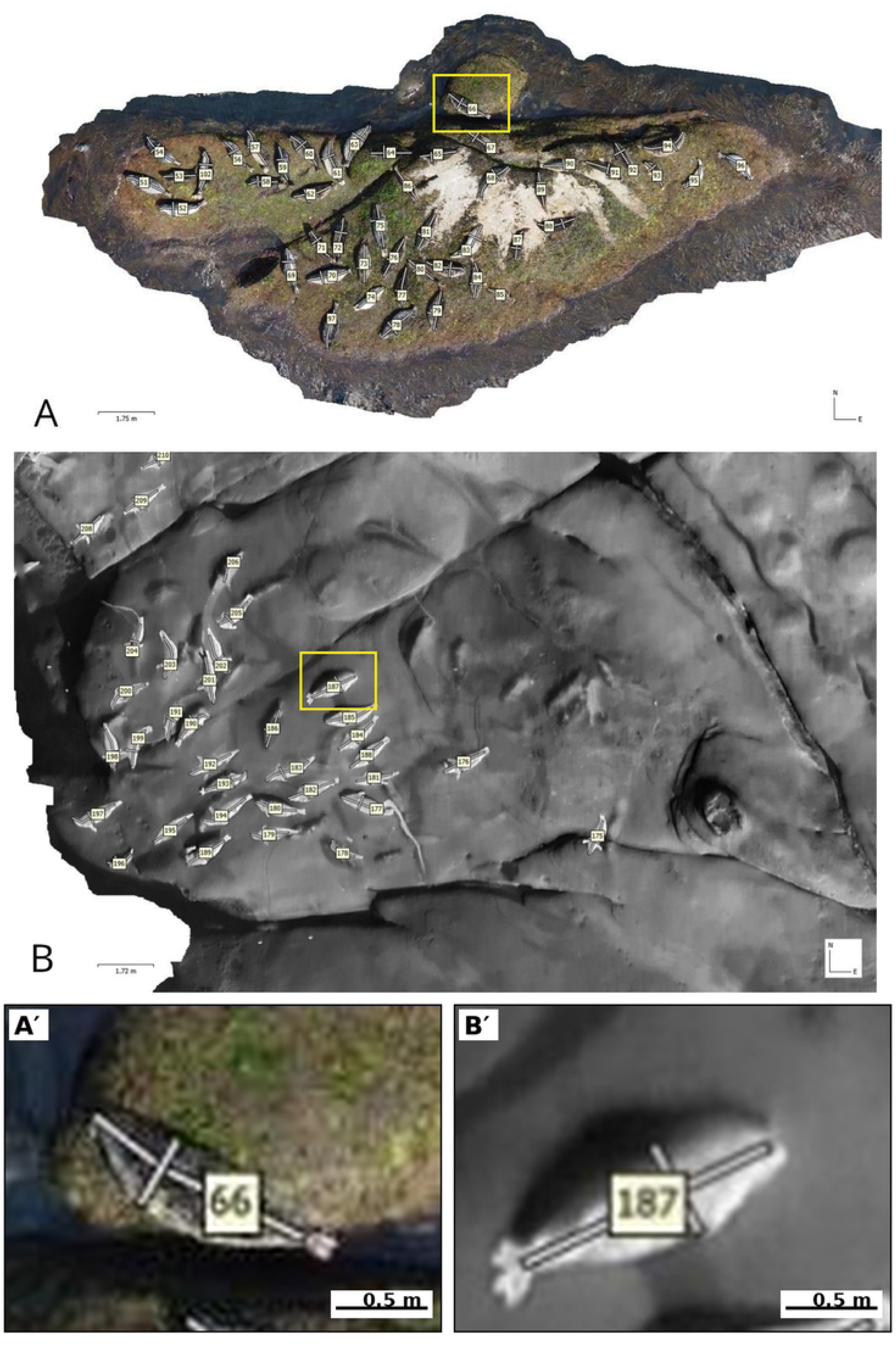
**Examples of manual annotation of *P.*** *vitulina* on (A) an RGB orthomosaic and (B) a thermal infrared (LWIR) orthomosaic (M3T, 20 m). Note the lower spatial resolution and blurred body boundaries in the thermal image compared with RGB. Both panels are reproduced at the same scale (scale bars 1.75 m and 1.72 m, respectively); insets A′ and B′ show a single annotated individual from each panel at a common, larger magnification, allowing the effective resolution of the two sensors to be compared directly.

The pipeline comprised three DeepLabV3+ segmentation models: (1) a water-removal model, (2) a first-level segmentation model for colony-level seal-versus-background classification, and (3) a second-level segmentation model for individual body-contour refinement. The first model masked water pixels in raw images before orthomosaic construction. The second model performed binary segmentation on the tiled orthomosaic. The third model refined the mask of each individual on a crop extracted from the bounding box of the second model’s prediction. Training sets were compiled separately for each model. The water-removal training set consisted of 800 image–mask pairs from haulouts with diverse substrates. The first-level segmentation training set comprised 4,101 pairs derived from 7 orthomosaics of spotted seals (*Phoca largha*) [11] and 10 harbor seal orthomosaics from this study. The second-level segmentation training set contained 2,757 pairs generated by automatic cropping with isolation of individual animals. For independent validation, a separate set of 31 orthomosaics (n = 2,483 manually annotated individuals) was reserved; none of these overlapped with the 10 *P. vitulina* or 7 *P. largha* training orthomosaics, and no individual animal appeared in both sets. The inclusion of cross-species (*P. largha*) training data was intended to increase morphological diversity.

#### 2.2.1. Calibration dataset and photogrammetric quality indicators

Orthomosaic geometric accuracy was quantified as the RMSE between the known length of each calibration rod and its manual measurement on the orthomosaic. To explain the variation in this error among surveys, five survey characteristics were examined: UAV platform, planned flight altitude, substrate type, GPS-anchoring density (ρ), and total camera location error (Camera Error Total, CET). The first three are categorical and fixed by survey design; the latter two are continuous and were taken from the Agisoft Metashape processing report.

GPS-anchoring density (ρ, cameras ha⁻¹) is the number of successfully aligned cameras per unit of orthomosaic coverage area:

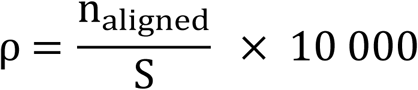

where n_aligned is the number of cameras aligned in Metashape and S is the orthomosaic coverage area (m²). The metric reflects image overlap and observation redundancy in the block adjustment: a higher camera density per unit area provides more connections between images and a more stable geometric solution.

Total camera location error (CET, m) is the root-mean-square discrepancy between the original GPS coordinates of the images and the coordinates of the same cameras after bundle adjustment:

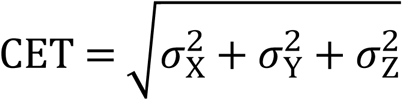

where σX, σY, and σZ are the root-mean-square deviations of the adjusted camera coordinates from the original GPS coordinates along the latitude, longitude, and elevation axes, respectively. CET is positional in form: it is the total camera location error of the Metashape ‘Average camera location error’ table, aggregating only the X, Y and Z components of the camera-position residuals, while orientation residuals (yaw, pitch, roll) and tie-point reprojection error are reported separately and are not part of it. It is integrative in content, however, because the adjusted camera positions are the end product of the entire bundle adjustment, so their residual against the recorded GPS positions responds to any factor that degrades that adjustment.

CET was treated as a candidate predictor of orthomosaic geometric accuracy for three reasons. First, it is a standard field of the Agisoft Metashape report and requires neither additional computation nor ground control points. Second, the factors it responds to span hardware (onboard GNSS accuracy, optics), flight geometry (pattern, overlap) and environment (substrate texture, illumination), so one number stands in for all of them. Third, unlike reprojection error or tie-point multiplicity, which are internal to the photogrammetric solution, CET is referenced to external absolute positioning, which makes it sensitive specifically to scale distortions in the orthomosaic.

UAV platform was a categorical factor with four levels (M3T, M2P, Mn3, Mn2). Planned flight altitude had three levels: 20 m (n = 42), 30 m (n = 2), and 40 m (n = 8). Substrate was classified into four types: slopes (n = 25), reefs (n = 7), boulders (n = 9), and plateaus (n = 6). Four surveys were left unclassified (No_data; S1 Table) because the terrain combined elements of several types and could not be assigned to a single class, and one seal survey was conducted on a sand beach (white_beach; n = 1).

### 2.3. Automated pipeline architecture

The automated pipeline consisted of eight sequential modules (Fig 4A). All three segmentation models were based on the DeepLabV3+ architecture with a ResNet-50 encoder pretrained on ImageNet and an Atrous Spatial Pyramid Pooling (ASPP) module with atrous rates of 6, 12, and 18 [28]. The choice of DeepLabV3+ was driven by three practical considerations. First, cascaded semantic segmentation with refinement can separate overlapping individuals with fewer training data than Mask R-CNN or YOLO-Seg, both of which require per-instance polygon annotation. Second, the ASPP module accommodates objects of different scales - from pups (∼75 cm) to adults (∼160 cm) - within a single input. Third, the computational requirements of DeepLabV3+ are lower than those of transformer-based architectures (ViT, SAM), allowing the model to be trained on hardware accessible to most field biologists. For first-level segmentation, the orthomosaic was divided into 256 × 256 pixel tiles with 25% overlap. The output layer was a binary mask with sigmoid activation. Dice loss [29–31], which optimizes overlap between predicted and reference masks, served as the loss function. Training was performed in R 4.3.2 with the tensorflow, keras3, and reticulate packages (TensorFlow 2.20 backend) on an NVIDIA RTX 5080 GPU. The architecture of the segmentation network itself is shown in Fig 4B.

**Fig 4.**
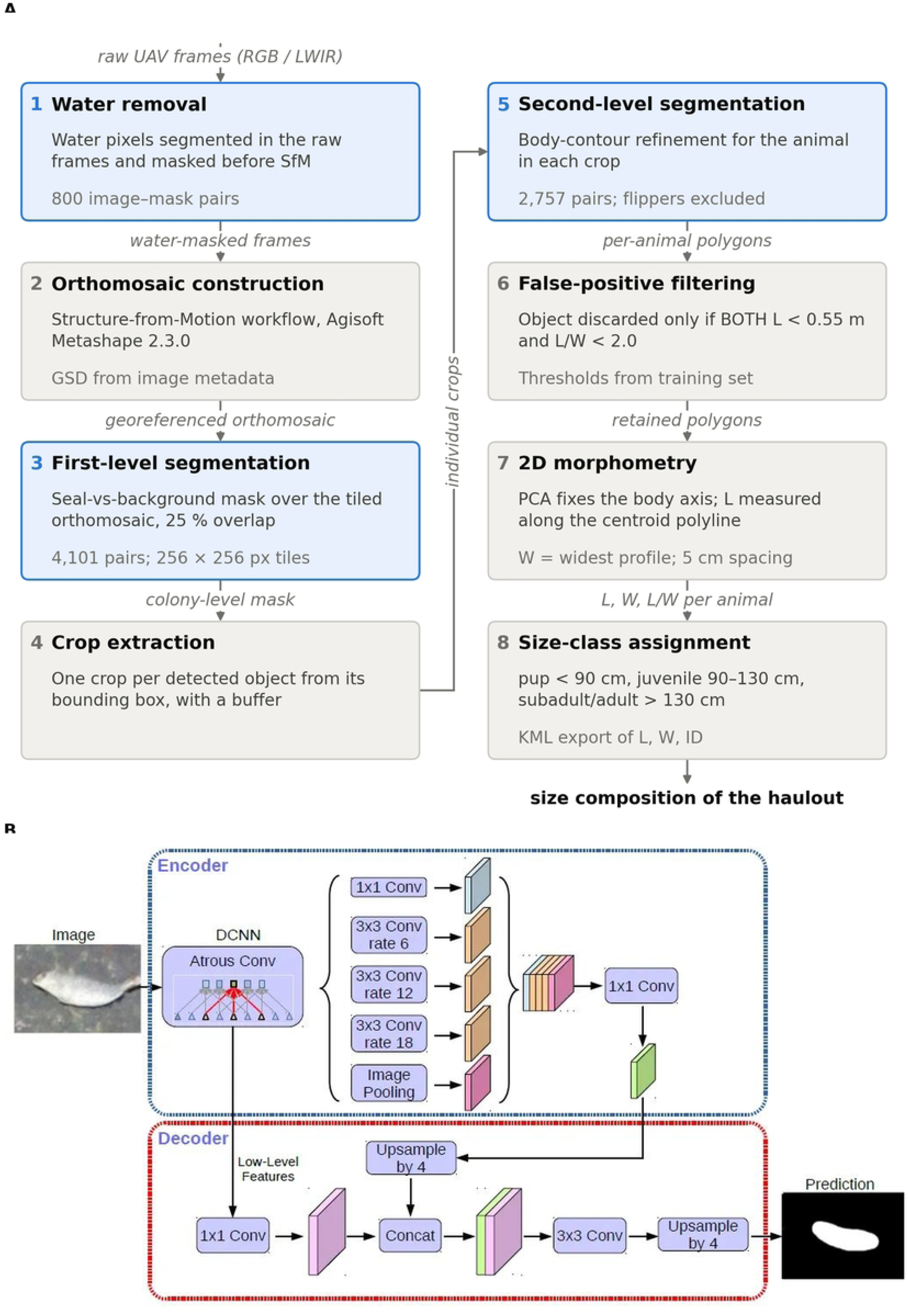
The automated pipeline. (A) The eight processing steps as a directed graph: node annotations give the key parameter of each step, italic labels on the edges name the data object passed to the next step, and the three steps shaded blue are the DeepLabV3+ instances of the cascade. (B) The DeepLabV3+ architecture shared by those three steps, with the ASPP module at atrous rates of 6, 12, and 18 (after [28]).

Training parameters were identical across all three models (see [32] for details): Adam optimizer; initial learning rate 1 × 10⁻⁴; ReduceLROnPlateau scheduler (patience = 5 epochs, factor = 0.2); early stopping on val_dice_coef (patience = 5 epochs); batch size 8; maximum 50 epochs. Augmentation included random flipping, rotation up to ±30°, and brightness and contrast variation of ±20%. The validation split was 20% (stratified).

### 2.4. False-positive filtering

After second-level masks were obtained, post-filtering was applied to remove artifacts. An object was excluded if two conditions were met simultaneously: length less than 0.55 m and length-to-width ratio less than 2.0. Both thresholds were determined from the training set, not from the validation data. The threshold of L ≥ 0.55 m was treated as the lower bound of neonatal *P. vitulina* body length. The filter was intentionally designed as a joint condition combining small size and atypical body shape. Small objects were not excluded by size alone, because they may correspond to pups whose accurate enumeration is a priority for demographic monitoring. Because these thresholds are species- and population-specific, they may require adjustment for species with different neonatal sizes or body proportions.

### 2.5. Morphometric parameter extraction

Two-dimensional measurements were extracted from binary polygons in the following sequence: (1) reprojection into the local UTM zone; (2) contour smoothing (smoothr package in R); (3) principal component analysis of the polygon vertices to establish body orientation only, the first principal component (PC1) defining the longitudinal direction and the second the transverse direction; PC1 is used only to orient the transverse-profile sampling, not as the measurement line; (4) construction of transverse profiles at 5 cm intervals along the principal axis; (5) derivation of body length (L) as the cumulative length of the polyline connecting the snout tip, the successive transverse-profile centroids, and the tail tip, so that L follows the curved body midline rather than the straight PC1 axis and thereby incorporates lateral body curvature; and (6) derivation of body width (W) as the maximum transverse width. W is a landmark-free maximum dorsal body width measured in dorsal projection, not an axillary girth or a circumferential measurement. W is thus the two-dimensional dorsal analogue of maximum girth, which is routinely recorded alongside axillary girth in standard pinniped morphometric protocols; an axillary position could not be anchored because the fore-flippers were deliberately excluded from the segmentation masks and therefore leave no landmark on the predicted polygon. The L/W ratio encoded body shape (Fig 5). L/W additionally served as a false-positive filter (objects with both length < 0.55 m and L/W < 2.0 were excluded; Section 2.4) but did not enter the size-class assignment, which is based on body length alone. Flippers were not included in the segmentation: the training masks delineated the body contour only (trunk and head), excluding both fore and hind flippers. Consequently, the predicted polygon did not encompass flippers, and the width measurement reflected the maximum dorsal body width rather than flipper span. The centroid polyline recovers curvature within the horizontal plane of the orthomosaic; dorso-ventral flexion is not observable in dorsal projection, so L remains a lower bound on true curvilinear body length for strongly arched postures. This limitation is intrinsic to two-dimensional photogrammetry and applies equally to the manual reference measurements, so it does not bias the automated-versus-manual comparison reported here.

**Fig 5.**
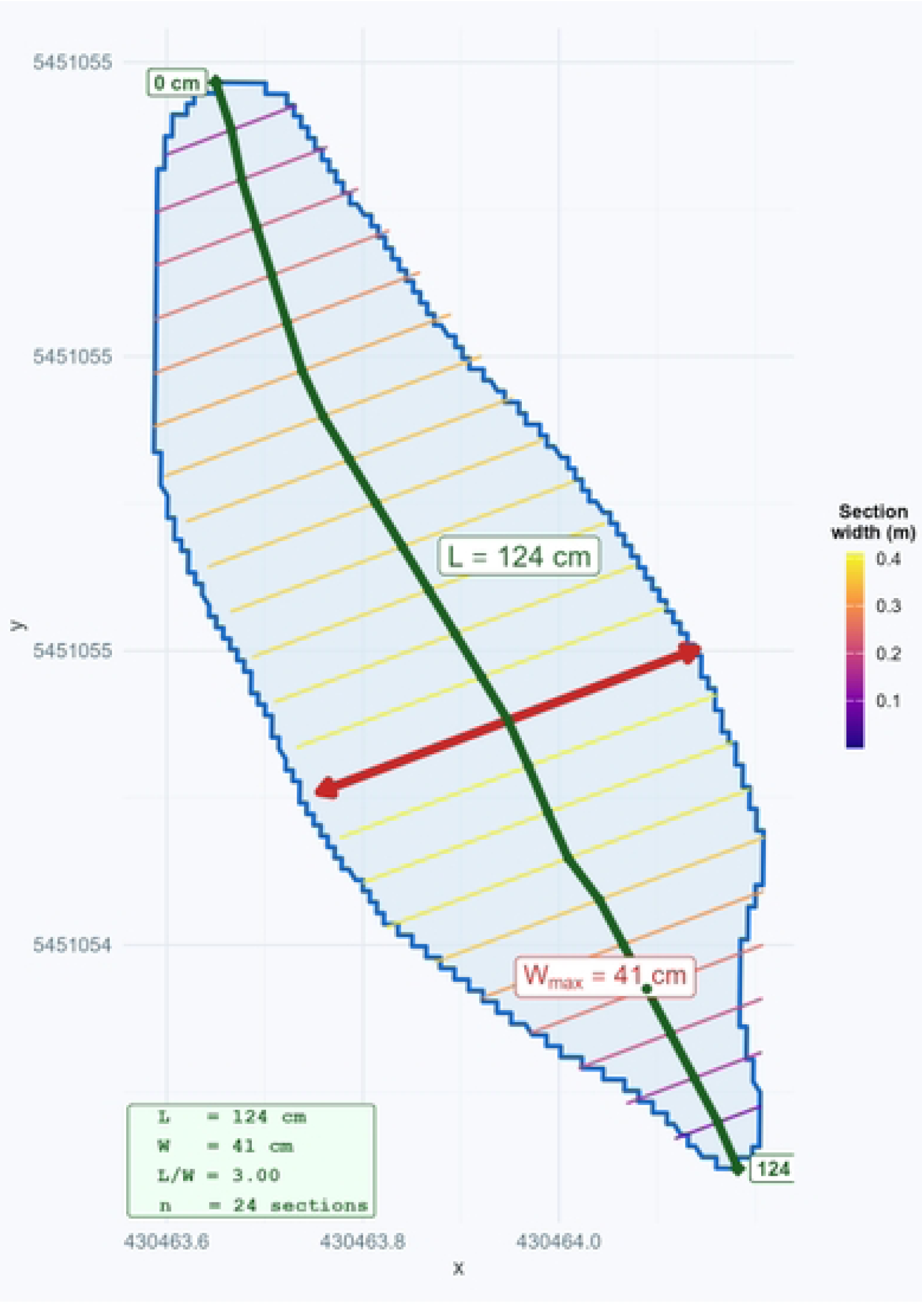
Schematic of the PCA-based method for extracting 2D morphometric parameters. The segmented seal polygon (blue) is oriented using the first principal component. Transverse profiles (colored lines, shaded by section width; see color scale) are constructed at 5 cm intervals. Body length (L, green) is measured along the midline connecting the snout tip, the transverse-profile centroids, and the tail tip, which follows body curvature; maximum width (W, red) is the widest profile.

### 2.6. Size-based classification

Based on published growth data for *P. vitulina* (see the supporting information and S2 Table), we adopted a three-class size system based on body length L: pups (L < 90 cm), corresponding to the lactation period and the first 1–2 months post-weaning; juveniles (90 ≤ L ≤ 130 cm), spanning approximately 2–3 months to ∼3 years of age; and subadults/adults (L > 130 cm), individuals approaching or having reached asymptotic length. The classification evaluates the pipeline’s ability to reproduce the size categories defined by the specified thresholds, using the labels pup, juvenile, and adult as convenient proxies for these size classes while acknowledging that biological age may not align perfectly with size-based boundaries due to individual variation in growth rate and to the spread of birth dates within the pupping season.

Because these classes are defined on length, both the reference and the predicted classes were assigned by applying the same two length cut-offs: the reference (“true”) classes to the lengths measured manually on the orthomosaic, and the predicted classes to the automatically measured lengths. No in vivo or visual age assessment was used. The task therefore measures how well the pipeline reproduces these length-defined size categories within the same information domain, the orthomosaic, in which the measurements are applied, rather than biological age as such; the labels pup, juvenile and subadult/adult serve only as convenient names for these size categories. Because of individual growth variability and the spread of birth dates within the pupping season, biological age does not map exactly onto the length boundaries (Section 4.4). Ninety-five percent confidence intervals for overall accuracy (OA) were estimated by the orthomosaic-level cluster bootstrap described in Section 2.7 (4,000 iterations).

### 2.7. Statistical analysis

Detection performance was evaluated using three standard metrics: Precision = TP/(TP+FP), Recall = TP/(TP+FN), and F1 = 2 × Precision × Recall / (Precision + Recall), where TP denotes true positives, FP false positives, and FN false negatives. A predicted polygon was scored as TP if its intersection-over-union (IoU) with the manual annotation reached ≥ 0.8. This threshold was chosen because subsequent morphometry requires close contour agreement, not merely correct localization.

Morphometric accuracy (body length, body width, calibration-rod length) was evaluated for TP objects using differences Δ = predicted − reference. Because manual annotations served as the reference for body length and width, these differences quantify the pipeline’s agreement with manual delineation on the same orthomosaic — its ability to reproduce expert annotation — rather than absolute accuracy against an animal’s true dimensions; only the calibration rods, whose lengths are known independently, provide an absolute (true-length) accuracy check. Any error common to the manual and automated measurements (e.g., posture, dorsal-projection distortion, or flipper exclusion) therefore cancels in Δ and is not captured by the body-size metrics. Four metrics were computed for each stratified subsample: bias (mean Δ), characterizing systematic error; mean absolute error (MAE), reflecting typical deviation magnitude; standard deviation (SD), characterizing random error; and root mean square error (RMSE), combining both components. Reporting bias and MAE jointly allows the error structure to be resolved: MAE ≈ |bias| indicates predominantly systematic error, whereas MAE >> |bias| indicates random noise dominance. The same error metrics were additionally computed in relative terms, as a percentage of the reference measurement.

Classification performance was evaluated by overall accuracy (OA), with 95% confidence intervals from the orthomosaic-level cluster bootstrap (4,000 iterations), together with the distribution of misclassifications across the three size classes. The robustness of the 90 and 130 cm class thresholds was assessed in a sensitivity analysis that shifted each threshold by ±5 and ±10 cm.

In order to assess the geometric accuracy of the orthomosaics, in addition to stratification by platform, substrate, and altitude, the continuous indicators ρ and CET were divided into terciles across the 22 calibration surveys. The tercile boundaries were 260 and 680 cameras ha⁻¹ for ρ, and 1.53 and 3.98 m for CET. Analysis of the factors affecting CET was performed on a combined dataset of 52 unique orthomosaics (the 22 calibration and 31 seal surveys, less the one orthomosaic [H0054] common to both). The monotonic association between each continuous indicator (rod length, ρ, and CET) and geometric error was quantified with Spearman’s rank correlation (r◻); because all four rod lengths occur in every orthomosaic, rod length varies within clusters and the 95% CI of its r◻ was obtained from the orthomosaic-level cluster bootstrap, whereas ρ and CET, which are fixed per orthomosaic, were correlated at the orthomosaic level (n = 22).

All detection, measurement, and classification metrics were stratified by sensor type, drone platform, substrate type, and flight altitude. Because animals located within the same orthomosaic are not statistically independent (they share a single photogrammetric reconstruction and therefore a common geometric error), all 95% confidence intervals for performance metrics (Precision, Recall, F1, OA, and the RMSE, bias, and MAE of body measurements) were obtained by a cluster bootstrap which resampled whole orthomosaics with replacement, rather than resampling individual animals, with 4,000 iterations. The same orthomosaic-level resampling was used to compare recall between RGB and thermal imagery, reported as the bootstrap 95% CI of the recall difference, in place of an animal-level χ² test that would treat clustered detections as independent. Platform, substrate, and flight-altitude effects on the geometric (calibration-rod) error were screened with the Kruskal–Wallis test, with pairwise Mann–Whitney tests as the post hoc procedure; the same omnibus test was applied across CET terciles and to the factors affecting CET. For all statistical tests, the significance level was set at α = 0.05. To control the family-wise error rate, pairwise Mann–Whitney comparisons were performed only following a significant Kruskal–Wallis omnibus test, and the resulting p-values were Bonferroni-adjusted within each such family. No global correction was applied across the full set of stratified analyses (by sensor, platform, substrate, and altitude); because these strata were examined to characterize where and how error varies rather than to test a single pre-registered hypothesis, the stratified comparisons are interpreted as exploratory, and the corresponding p-values should be read as descriptive rather than confirmatory. All analyses were performed in R 4.3.2.

The per-animal morphometric measurements and the calibration-rod measurements underlying every analysis reported here are provided in S1 Dataset and S2 Dataset. Trained DeepLabV3+ models, R scripts for morphometric extraction and pipeline configuration files are deposited at https://github.com/ivan-npwc/PV_PL.

### 2.8. Ethics statement

All drone surveys were conducted at altitudes ≥ 20 m, minimizing behavioral disturbance to *P. vitulina* [33]. Flights complied with Transport Canada regulations for drone operations in wildlife areas. No physical contact with animals occurred; all data were obtained by remote sensing. Because the study involved exclusively non-invasive remote sensing from altitudes ≥ 20 m with no animal capture, handling, or disturbance beyond the presence of the aircraft, no IACUC or animal ethics committee approval was required under applicable Canadian and U.S. regulations. Field operations were conducted under Fisheries and Oceans Canada marine mammal licence XMM 1 2021. The species is not listed under the Species at Risk Act, so no permit under that Act was required.

## 3. Results

### 3.1. Segmentation model training

All three DeepLabV3+ models were trained for 50 epochs on an NVIDIA RTX 5080 GPU. The water-removal model reached a best validation Dice of 0.960 at epoch 28 (early stopping at epoch 38) with 800 training pairs. The first-level model reached Dice = 0.911 at epoch 46 of 50 (4,101 pairs). The second-level model reached Dice = 0.939 at epoch 48 of 50 (2,757 pairs). Learning curves (S1–S3 Figs) showed monotonic decrease in Dice loss with no train–validation divergence, indicating no overfitting. The ReduceLROnPlateau scheduler reduced the learning rate five-fold upon stagnation: once for the second-level model, three times each for the water-removal and first-level models.

### 3.2. Detection accuracy

The pipeline was evaluated on 2,483 manually annotated *P. vitulina*. It correctly identified 2,298 individuals (TP), missed 185 (FN), and generated 247 false positives (FP), yielding Precision = 0.903 (95% CI 0.879–0.923), Recall = 0.925 (0.891–0.958), and F1 = 0.914 (0.891–0.936). Of the 185 FN, 10 (5.4%) were attributable to insufficient polygon overlap; the remaining 175 were genuine detection failures.

#### 3.2.1. Sensor type: RGB versus thermal infrared

Sensor modality had the largest effect on detection performance among all factors examined. RGB imagery (n = 1,821) yielded F1 = 0.928 (95% CI 0.908–0.943) with high recall (0.957; 0.940–0.971), indicating that animals were rarely missed in color images. Thermal infrared imagery (n = 662) produced a lower F1 = 0.874 (0.848–0.920), driven by reduced recall (0.838; 0.810–0.903 vs. 0.957) rather than precision (0.913 vs. 0.900). The RGB–thermal recall difference was 11.9 percentage points (cluster-bootstrap 95% CI 5.3–15.2), so the advantage of RGB for detection remained clearly supported once within-orthomosaic dependence was accounted for. Thermal data contained 107 FN compared with 78 in the 2.8-times-larger RGB subset. In dense aggregations, individual thermal signatures merge into continuous heat fields, causing systematic segmentation failure for adjacent individuals.

### 3.3. Geometric accuracy of orthomosaics (RGB only)

Aggregate geometric metrics, computed on all calibration-rod measurements (n = 545), were: bias = −1.22 cm (95% CI −3.23 to 0.61), MAE = 3.75 cm (2.32–5.52), SD = 6.56 cm (4.08–8.62), and RMSE = 6.67 cm (4.19–8.90; cluster bootstrap over orthomosaics). Per-survey RMSE varied widely, from 1.0 to 18.0 cm (S1 Table).

Absolute error increased monotonically with rod length, whereas relative error remained constant: mean relative MAE was 3.8–4.3% and mean relative RMSE 6.5–6.9% across the four rod lengths. Error pooled within each platform–altitude combination, over substrate, is shown against rod length in Fig 6. Because all four rod lengths were present in every orthomosaic, rod length varies within clusters, so its effect was assessed with Spearman correlations whose 95% confidence intervals were obtained by the orthomosaic-level cluster bootstrap. Absolute error was positively associated with rod length (Spearman’s r◻ = 0.27, cluster 95% CI 0.19–0.36), whereas relative error was not (r◻ = −0.02, cluster 95% CI −0.10 to 0.05).

**Fig 6.**
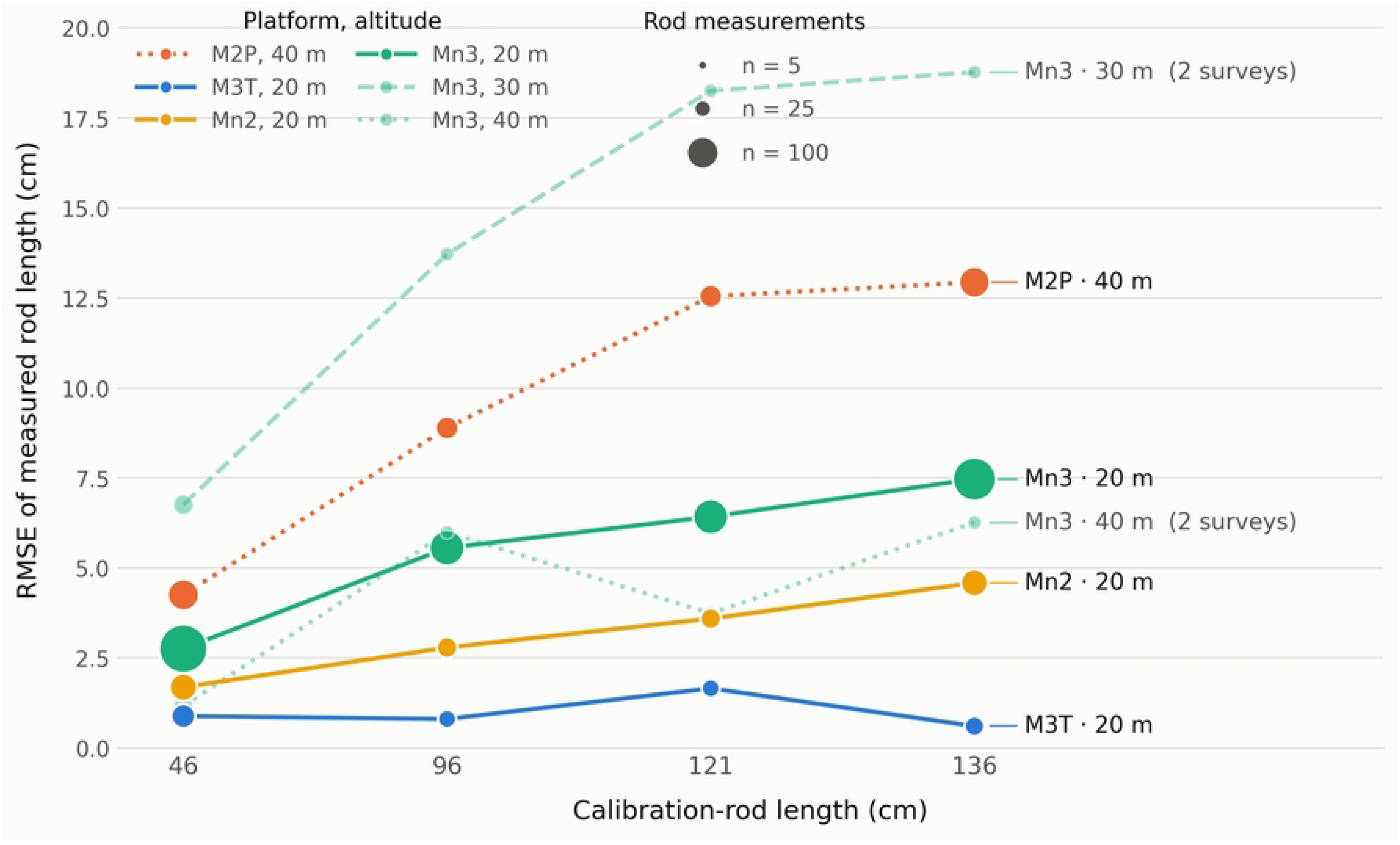
Geometric accuracy of calibration-rod measurements. Root-mean-square error (RMSE) of measured versus true rod length as a function of rod length, by platform (line color) and flight altitude (line style), pooled within each platform–altitude combination over the same 22 calibration orthomosaics used in Section 3.3. Marker area is proportional to the number of rod measurements behind each point. Under-sampled combinations (median n < 8 rods per rod length) are drawn faded; every series is labelled at the right-hand end of its line, and the number of orthomosaics behind each faded series is given with its label.

Platform, flight altitude, and substrate were analyzed jointly because their effects are confounded. At the orthomosaic level (n = 22 orthomosaics, the appropriate unit), only flight altitude was significantly associated with geometric error (Kruskal–Wallis H = 6.60, p = 0.037); platform (H = 6.91, p = 0.075) and substrate (H = 6.27, p = 0.18) were not significant once calibration rods were no longer treated as independent observations. The lowest error was obtained with M3T on a plateau at 20 m (RMSE ≈ 1.0 cm) and the highest reliable error with M2P on rocky slopes at 40 m (RMSE ≈ 10.0 cm). The single highest value (Mn3 on reefs at 30 m, ≈ 18 cm) rested on only 12 rod measurements. Post hoc comparisons indicated that this altitude effect was driven by the M3T–20 m–plateau combination; once it was excluded, the differences became non-significant. The absence of a significant substrate term should not be read as evidence that substrate is unimportant. With 22 orthomosaics distributed unevenly among four substrate classes, and with substrate largely confounded with platform and altitude, the orthomosaic-level test has little power to isolate it. Calibration rods were laid directly on the substrate surface and were not levelled (Section 2.1), so on inclined ground part of the true rod length lies outside the horizontal plane and is foreshortened in the orthomosaic; this acts to inflate apparent error on slopes and therefore biases the test towards, not away from, detecting a substrate effect. The exploratory pattern reported in Section 3.6, in which flat substrates outperform slopes for both detection and classification, is consistent with a substrate effect that is present but under-resolved at n = 22 orthomosaics.

The relationship between GPS-anchoring density (ρ) and accuracy was weak and non-monotonic: RMSE was 6.70 cm (cluster-bootstrap 95% CI 3.12–9.32) in the low-ρ tercile, 7.94 cm (1.71–11.97) in the middle tercile, and 5.31 cm (2.52–9.49) in the high-ρ tercile (at the orthomosaic level, density was not associated with RMSE: Spearman’s r◻ = −0.15, p = 0.50).

Tercile stratification by CET yielded a monotonic relationship. The low tercile (CET ≤ 1.53 m) had RMSE = 2.31 cm (cluster-bootstrap 95% CI 1.44–3.14), the middle tercile (1.53–3.98 m) RMSE = 8.07 cm (2.85–11.90), and the high tercile (> 3.98 m) RMSE = 9.35 cm (5.62–12.53). At the orthomosaic level (n = 22 orthomosaics, the appropriate unit of analysis), CET was strongly associated with geometric RMSE (Spearman’s r◻ = 0.78, p < 0.001; Kruskal–Wallis across terciles H = 9.87, p = 0.007); because each tercile contains only 7–8 orthomosaics, pairwise tercile comparisons should be regarded as exploratory.

The only robust predictor of CET was flight altitude (H = 13.4, p = 0.001; n = 52). Median CET was 1.24 m at 20 m, 7.87 m at 30 m, and 5.25 m at 40 m. The platform effect disappeared once altitude was controlled for (H = 2.3, p = 0.31), and substrate was non-significant (H = 8.4, p = 0.08).

### 3.4. Agreement of automated and manual body measurements (RGB and thermal)

Automated body-length measurements for 2,298 TP detections yielded aggregate RMSE = 16.1 cm (bias = −5.3 cm, MAE = 12.4 cm, relative error = −5.04%) (cluster-bootstrap 95% CIs over orthomosaics: RMSE 13.0–19.1, bias −10.1 to −1.0, MAE 10.0–15.1 cm). Width measurements (n = 2,102) agreed more closely with manual annotation: RMSE = 5.3 cm (bias = −1.2 cm, MAE = 3.2 cm, relative error = −3.75%) (95% CIs: RMSE 4.2–6.3, bias −1.8 to −0.7, MAE 2.8–3.5 cm), reflecting clearer lateral body contours. Length and width were measured independently for each individual; consequently, some individuals with a valid length lacked a valid width (e.g., where a reliable maximum width could not be determined from the body contour), and width was therefore assessed on 196 fewer detections than length (n = 2,102 vs. 2,298 for length). This reflects independent per-axis measurement rather than any additional filtering of detections.

#### 3.4.1. RGB versus thermal

Stratification by sensor revealed that aggregate error was driven largely by thermal data (Figs 7 and 8). For RGB (n = 1,743), the pipeline achieved RMSE = 12.5 cm with near-zero bias (−0.6 cm) and MAE = 9.7 cm (cluster-bootstrap 95% CIs: RMSE 11.8–13.4, bias −2.2 to 0.7, MAE 9.1–10.4 cm). The relationship MAE >> |bias| (9.7 >> 0.6 cm) indicates predominantly random errors: over- and underestimates cancel at the population level, but individual deviations average ∼10 cm. The body-length error showed no systematic dependence on manually measured body length (Fig 9).

**Fig 7.**
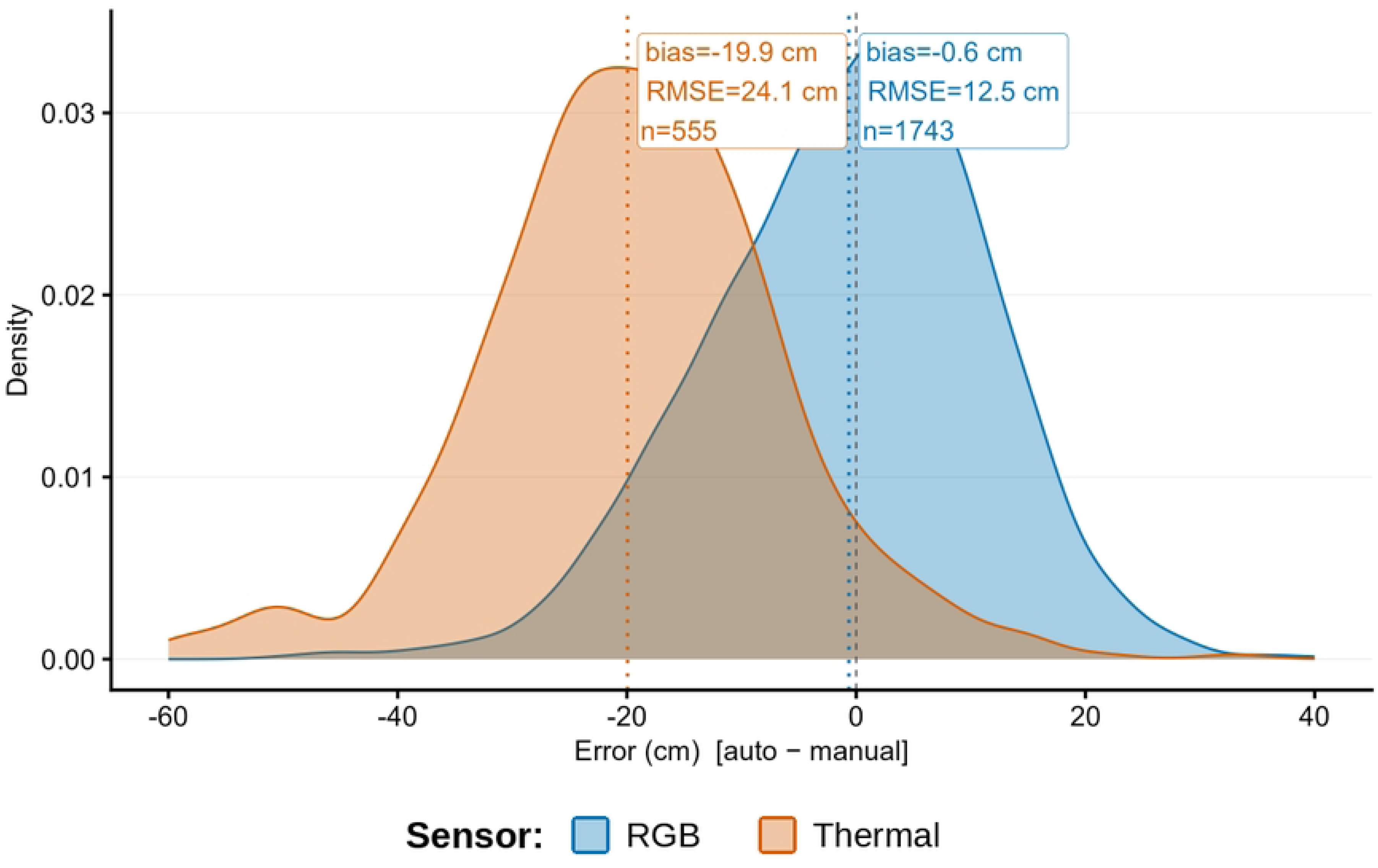
Distribution of automated body-length measurement errors (ΔL = automated − manual, cm) by sensor type. Automated RGB lengths show near-zero bias relative to manual annotation (−0.6 cm, RMSE = 12.5 cm), whereas automated thermal lengths are systematically shorter (−19.9 cm, RMSE = 24.1 cm).

**Fig 8.**
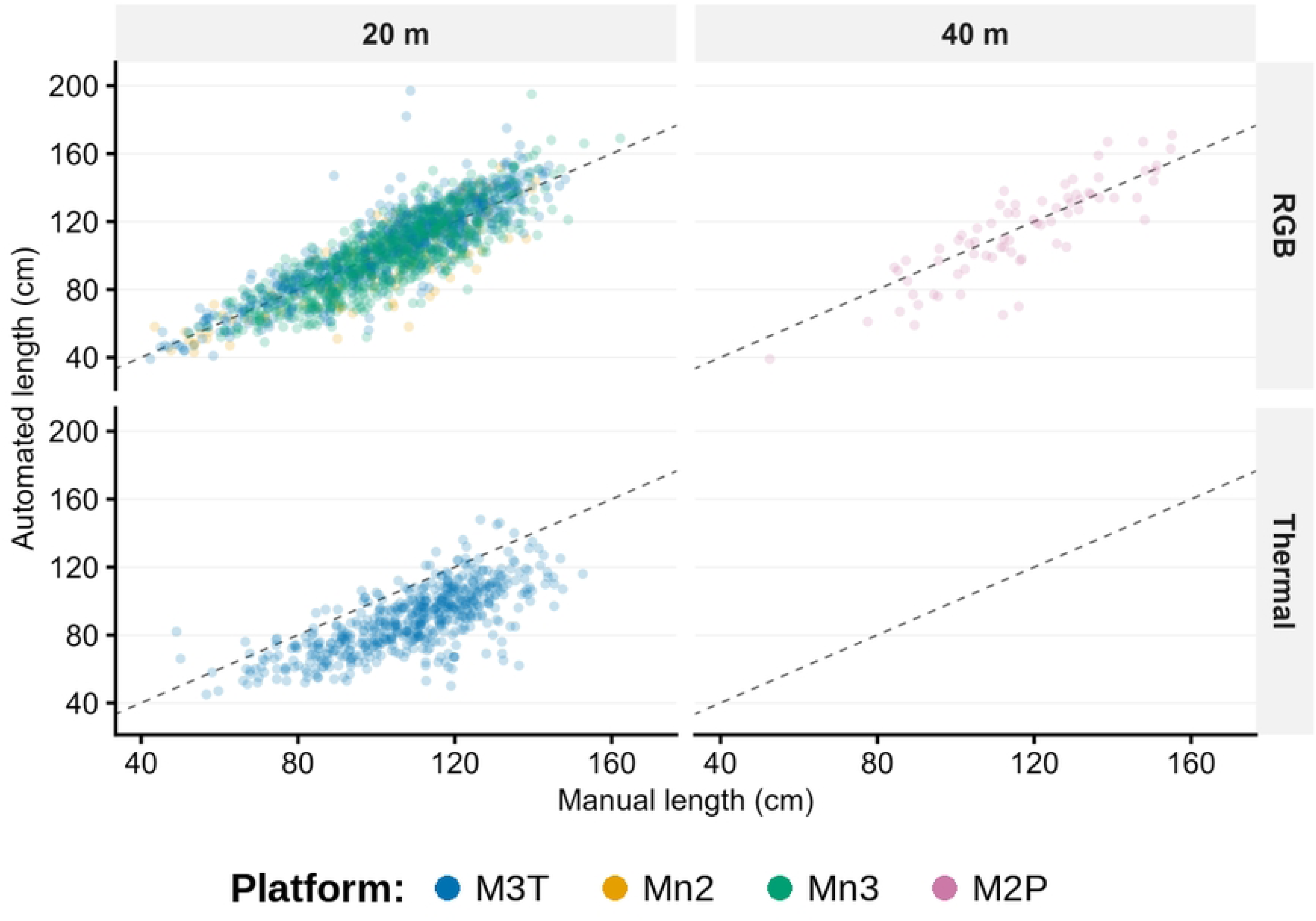
Automated versus manual body-length measurements, stratified by sensor type, with the 1:1 line. Panels are arranged by flight altitude (columns: 20 m and 40 m) and sensor (rows: RGB and thermal), with points colored by platform (M3T, Mn2, Mn3, M2P); the thermal / 40 m panel is empty because no thermal surveys were flown at 40 m.

**Fig 9.**
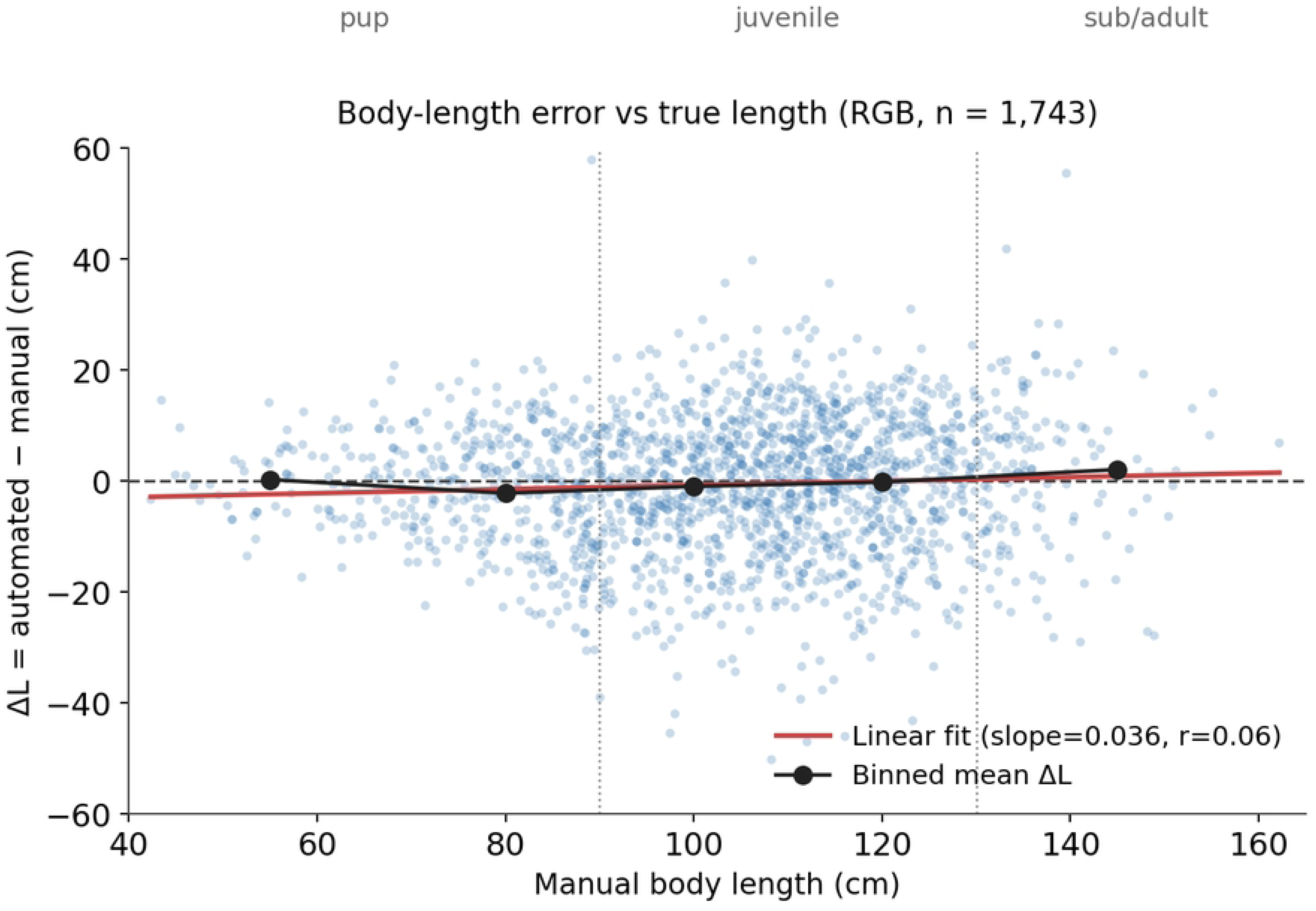
Automated body-length measurement error (*ΔL* = automated − manual, cm) as a function of manually measured body length, for RGB detections (n = 1,743). The dashed line marks zero error; the red line is a linear fit (slope = 0.04, r = 0.06) and the black line connects binned means. Dotted vertical lines mark the 90 and 130 cm size-class boundaries. Error is centred near zero across the whole length range, indicating that the random measurement error is approximately independent of body size.

For thermal imagery (n = 555), RMSE = 24.1 cm, bias = −19.9 cm, MAE = 20.8 cm (∼19% of the mean body length) (cluster-bootstrap 95% CIs: RMSE 22.3–25.1, bias −20.7 to −18.7, MAE 19.4–21.7 cm). The closeness of MAE to |bias| (20.8 ≈ 19.9 cm) shows almost entirely systematic error - nearly all measurements were uniformly underestimated. On the same platform (M3T, 20 m), thermal bias differed from RGB by 22.2 cm (cluster-bootstrap 95% CI 20.3–23.3, resampling 7 RGB vs. 5 thermal orthomosaics rather than individual animals). This bias is reproducible and substrate-independent; because both the manual reference and the automated length are measured on the same LWIR orthomosaic, it does not represent an absolute sensor error but a systematic segmentation–annotation difference at the blurred, low-contrast body margins (mechanism detailed in Section 4.3). This loss of edge definition is directly visible in the thermal orthomosaic of Fig 3B, where body outlines are blurred relative to the RGB image in Fig 3A. Width measurements confirmed the same pattern: RGB MAE = 2.9 cm with bias −0.7 cm (random error); thermal MAE = 4.0 cm with bias −2.9 cm (systematic underestimation) (Fig 8).

### 3.5. Automated size-based classification

The “true” class of each individual is the size category based on manually measured length: pups (< 90 cm), juveniles (90–130 cm), subadults/adults (> 130 cm). We evaluate how accurately automated measurements reproduce this categorization. All classification errors are direct consequences of length-measurement errors. Analysis is restricted to RGB data (n = 1,743).

#### 3.5.1. Overall accuracy

Threshold classification achieved OA = 78.7% (95% CI 76.4–81.0%, cluster bootstrap over orthomosaics). Of 371 misclassified individuals, most were juveniles assigned to pups (147, mean underestimation −18.2 cm) or to subadults/adults (120, mean overestimation +15.4 cm).

The subadult/adult class lost 27% of its members to the juvenile class due to length underestimation. A sensitivity analysis showed that the 90 and 130 cm thresholds are close to optimal for reproducing the manual size categories: shifting them by ±5 cm reduced overall accuracy by 1–3 percentage points, and by ±10 cm by 4–9 points.

#### 3.5.2. Classification uncertainty zones at size-class boundaries

The geometric accuracy of an orthomosaic (RMSE on the calibration rods) determines the classification uncertainty zones at the size-class boundaries. Each zone is defined as ±2 × RMSE around the corresponding boundary (90 and 130 cm). Because CET is the most robust predictor of orthomosaic geometric accuracy (orthomosaic-level Spearman’s r◻ = 0.78, p < 0.001; Section 3.3), the uncertainty zones were stratified by CET tercile, which links the photogrammetric reconstruction quality of a given orthomosaic to the expected reliability of classification.

The stratification results are presented in Table 1.

**Table 1.** Classification uncertainty zones by CET tercile.

| CET tercile | Geometric RMSE (cm) | n | Pup/juvenile zone (cm) | Juvenile/subadult zone (cm) | % in zones | Overall OA | OA within zones | OA outside zones |
| --- | --- | --- | --- | --- | --- | --- | --- | --- |
| Low ($\leq 1.53$ m) | 2.31 | 1157 | 85–95 | 125–135 | 19.6% | 78.6% | 55.9% | 84.1% |
| Middle (1.53–3.98 m) | 8.07 | 467 | 74–106 | 114–146 | 71.5% | 78.8% | 71.9% | 96.2% |
| High ($> 3.98$ m) | 9.35 | 119 | 71–109 | 111–149 | 79.8% | 79.8% | 76.8% | 91.7% |
| **Aggregate** | **6.67** | **1743** | **77–103** | **117–143** | **61.6%** | **78.7%** | **71.1%** | **90.9%** |

*Uncertainty zones: class boundary $\pm 2 \times$ RMSE for the 90 and 130 cm boundaries. Geometric RMSE is the value measured on the calibration rods within the corresponding CET tercile. OA, overall accuracy of the threshold classifier. The number of orthomosaics underlying each tercile is small (low, 19; middle, 4; high, 3 orthomosaics); per-tercile estimates are therefore exploratory and should be interpreted with caution.*

At CET ≤ 1.53 m (low tercile; geometric RMSE = 2.31 cm; n = 1157 individuals, 66.4% of the full sample), the uncertainty zones are narrow: 85–95 cm at the pup/juvenile boundary and 125–135 cm at the juvenile/subadult boundary. Only 19.6% of individuals fall within these zones.

At CET 1.53–3.98 m (middle tercile; RMSE = 8.07 cm; n = 467), the zones widen to 74–106 and 114–146 cm, encompassing 71.5% of individuals.

At CET > 3.98 m (high tercile; RMSE = 9.35 cm; n = 119), the zones span 71–109 and 111–149 cm, including 79.8% of individuals.

### 3.6. Effects of survey conditions across the pipeline

The same platform–sensor–substrate–altitude stratification was applied to all three tasks (Table 2). Because these factors are constant within an orthomosaic, the factor-level tests reported below treat animals as non-independent and are therefore exploratory (Section 2.7); the consistent direction of each effect across detection, measurement, and classification is more informative than any single test. In Table 2, n (TP) is the number of true-positive detections and n (rods) the number of calibration-rod measurements; geometric and body-length error are both reported as RMSE with MAE in parentheses (cm), with body-length error given RGB-only for comparability (thermal is shown separately under Sensor). Geometric RMSE is the calibration-rod error pooled within each level over the n (rods) shown; it is available for RGB orthomosaics only and is reported for sensor, platform, and altitude but not substrate, whose rod coverage is confounded with platform and altitude (the orthomosaic-level estimate and confidence intervals are given in Section 3.3).

**Table 2.** Effect of survey conditions on geometric accuracy, detection, body measurement, and size classification. Metric definitions and caveats are given in Section 3.6; the per-combination detection breakdown is in S3 Table.

| Factor | Level | n (TP) | Detection F1 | n (rods) | Geometric RMSE (MAE) cm | Body-length RMSE (MAE) cm | Classification OA (%) |
| --- | --- | --- | --- | --- | --- | --- | --- |
| Sensor | RGB | 1,743 | 0.928 | 545 | 6.7 (3.8) | 12.5 (9.7) | 78.7 |
|  | Thermal | 555 | 0.874 | — | — | 24.1 (20.8) | — |
| Platform | M3T (RGB) | 646 | 0.955 | 52 | 1.0 (0.7) | 11.9 (8.6) | 80.7 |
|  | M2P (40 m) | 72 | 0.960 | 107 | 10.0 (6.8) | 14.9 (11.5) | 75.0 |
|  | Mn3 | 871 | 0.908 | 305 | 6.4 (3.5) | 12.5 (10.1) | 78.5 |
|  | Mn2 | 154 | 0.914 | 81 | 3.4 (2.5) | 14.1 (10.9) | 73.4 |
| Substrate | Plateaus | 192 | 0.962 | — | — | 12.5 (8.4) | 81.8 |
|  | Boulders | 435 | 0.914 | — | — | 11.2 (9.0) | 82.1 |
|  | Slopes | 942 | 0.938 | — | — | 13.0 (10.1) | 76.5 |
|  | Reefs | 120 | 0.863 | — | — | 12.9 (10.3) | 80.0 |
|  | White beach | 54 | 0.893 | — | — | 12.8 (10.4) | 75.9 |
| Altitude | 20 m | 1,671 | 0.926 | 391 | 4.8 (2.6) | 12.4 (9.6) | 78.9 |
|  | 40 m | 72 | 0.960 | 130 | 9.3 (6.1) | 14.9 (11.5) | 75.0 |

#### Sensor

RGB outperformed thermal on every metric: detection F1 was 0.928 versus 0.874, and RGB body-length MAE was 9.7 cm versus 20.8 cm for thermal. The thermal deficit was a systematic underestimation independent of substrate (Section 3.4.1), confirming that thermal imagery supports detection but not quantitative morphometry.

#### Platform

The M3T gave the most accurate morphometry (RGB MAE 8.6 cm) and the highest classification accuracy (OA 80.7%); the lightweight Mini platforms were comparable for detection but less accurate for measurement and classification (Mn3 OA 78.5%, Mn2 73.4%).

The opposite signs of platform bias (M3T +1.0 to +3.0 cm; Mini platforms −0.7 to −3.5 cm, the largest values occurring on slopes) point to different error sources: boundary inclusion at fine ground sample distance versus edge-pixel loss at coarser resolution (Fig 10). The M2P was flown only at 40 m, confounding platform with altitude.

**Fig 10.**
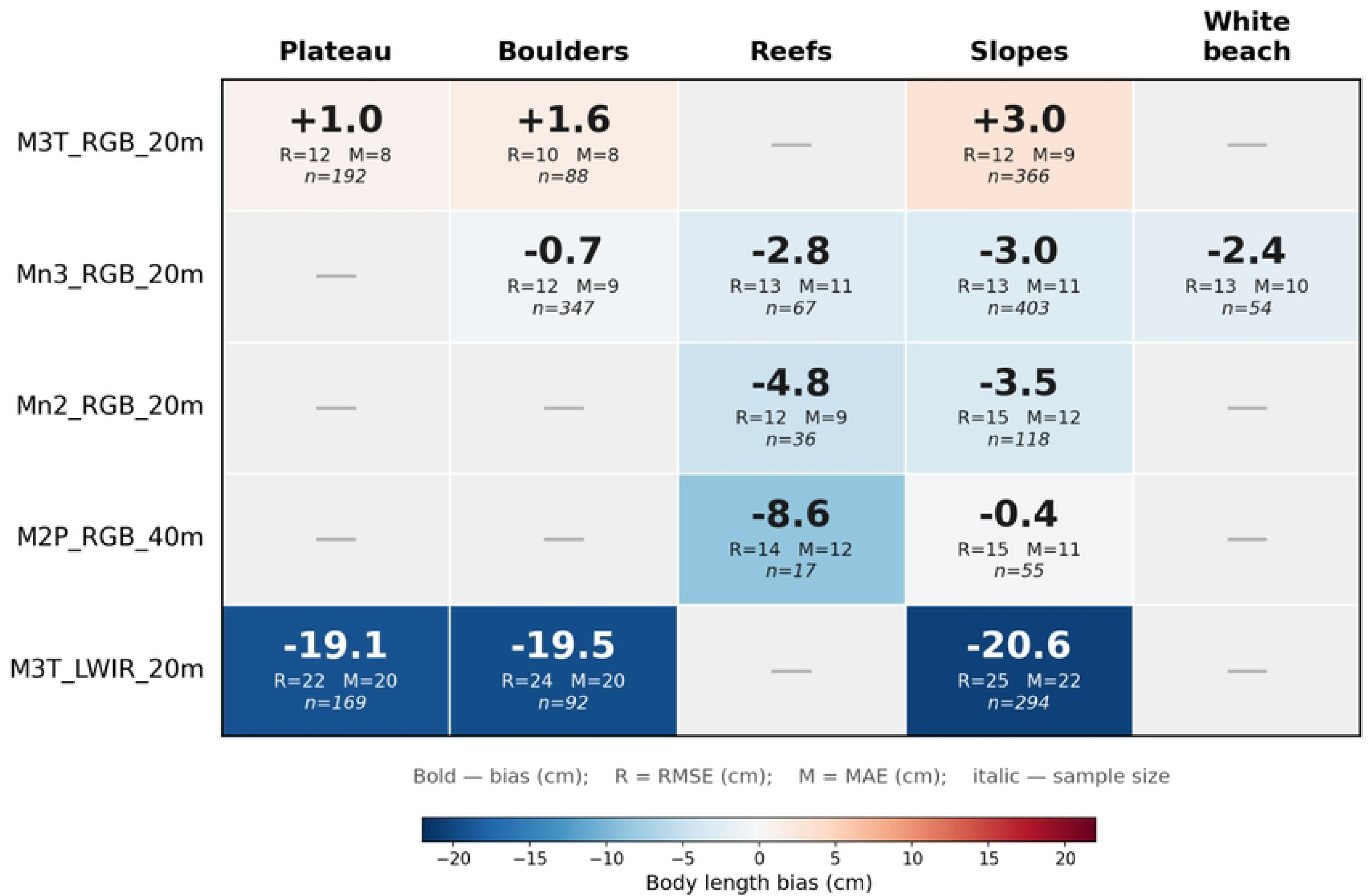
Heatmap of automated body-length measurement bias (cm) by platform–sensor–substrate combination. Bold: bias (cm); R: RMSE (cm); M: MAE (cm); italic: sample size. Positive bias (warm colors): systematic overestimation; negative bias (cool colors): underestimation. Gray cells: untested combinations.

#### Substrate

Flat, homogeneous surfaces performed best across tasks: plateaus yielded the highest detection F1 (0.962), and plateaus and boulders the highest classification accuracy (OA 81.8% and 82.1%) and the lowest measurement error, whereas slopes were the least accurate (OA 76.5%), where surface inclination distorts the dorsal projection of the body (Fig 10).

#### Altitude

Lower flight altitude improved every metric: at 20 m, classification accuracy was 78.9% and RGB body-length MAE 9.6 cm, declining to 75.0% and 11.5 cm at 40 m, where a coarser ground sample distance increases geometric error.

#### Geometric quality

Overall classification accuracy did not vary with orthomosaic geometric quality (OA 78.6%, 78.8%, and 79.8% across the low, middle, and high CET terciles), confirming that errors are driven by per-individual body length measurement error rather than by reconstruction accuracy. What the geometry does control is the width of the uncertainty zones around the 90 and 130 cm boundaries (±2 × calibration-rod RMSE; Section 3.5, Table 1): at CET ≤ 1.53 m the rod RMSE is only 2.31 cm and just ∼20% of individuals fall in the borderline zones (84.1% accuracy among the ∼80% outside them), whereas at CET > 1.53 m the rod RMSE rises to 8–9 cm and the zones widen to encompass more than 70% of the sample. In practice, under good photogrammetric conditions (CET ≤ 1.53 m) only individuals whose automated length falls in the 85–95 or 125–135 cm ranges need be flagged as “borderline” for visual verification; under poorer conditions (CET > 1.53 m) only individuals shorter than ∼77 cm or longer than ∼143 cm can be classified with high confidence.

## 4. Discussion

### 4.1. Pipeline performance in the context of published data

The cascaded DeepLabV3+ pipeline achieved F1 = 0.914 on 2,483 independently annotated seals, placing it in the upper quartile of published pinniped detection results. Infantes et al. [6] reported F1 = 0.95–0.97 for *P. vitulina* on homogeneous substrate with a single platform. Carroll et al. [9] achieved F1 = 0.953 on rocky substrates. Dujon et al. [7] showed strong dependence on morphology, density, and substrate across three marine taxa (recall 0.75–0.94; precision 0.20–0.74). For Australian fur seals (*Arctocephalus pusillus*) on heterogeneous substrate, recall was 0.79 with low precision (0.27) due to boulder-triggered false positives. Liu et al. [34] proposed a hierarchical YOLOv10/YOLOv7 scheme for spotted seals achieving precision = 0.942.

Our result is notable because it was obtained on a substantially more heterogeneous dataset - four platforms, two sensors, five substrates - conditions under which most published pipelines have not been tested. Direct comparison of F1 values across studies is complicated by differing TP definitions: we used IoU ≥ 0.8 (contour accuracy is critical for morphometry), whereas most studies apply IoU ≥ 0.5 or centroid matching. Relaxing our threshold to IoU ≥ 0.5 would likely push aggregate F1 above 0.93–0.94, comparable to Infantes et al. [6] and Carroll et al. [9] on less heterogeneous data.

The two-level scheme confirmed the cascaded architecture: single-level segmentation (Dice = 0.911) produced merged polygons in dense aggregations, which the cascade eliminated. A similar hierarchical strategy was independently proposed for spotted seals [34], supporting the generality of this approach. In the initial phase, Trukhanova et al. [11] surveyed spotted seal (*P. largha*) haulouts in Karaginsky Gulf (southwestern Bering Sea) using manual measurements of 4,057 individuals across 64 drone flights. That study confirmed the practical value of UAV photogrammetry for pinniped abundance and size composition assessment but relied entirely on manual processing without calibration references, precluding quantitative error characterization by platform or altitude. The cascaded DeepLabV3+ pipeline presented here addresses these limitations by automating detection, measurement, and classification while providing stratified accuracy estimates across survey conditions.

### 4.2. Geometric accuracy of orthomosaics and the CET predictor

CET responds to hardware, geometric, and environmental factors alike, which explains its position as the most robust predictor of geometric accuracy (orthomosaic-level Spearman’s r◻ = 0.78, p < 0.001), whereas platform, altitude, and substrate show confounded effects and GPS-anchoring density (ρ) shows only a non-monotonic relationship. The only robust predictor of CET itself is flight altitude (H = 13.4, p = 0.001); the platform and substrate effects do not persist once altitude is controlled for (H = 2.3, p = 0.31).

Image overlap is a necessary but not sufficient condition for accuracy, which also depends on the quality of the GNSS solution, the camera optics, scene texture, and the accuracy of the lens-distortion model. CET, which integrates all of these factors, proved a considerably stronger predictor than ρ (orthomosaic-level Spearman’s r◻ = 0.78 versus −0.15).

A CET ≤ 1.53 m yields a relative RMSE of about 2%, which for *P. vitulina* (∼150 cm) corresponds to an error on the order of 3 cm. At CET > 1.53 m the error rises to 6–10%, and at CET > 3.98 m size-class classification becomes unreliable.

### 4.3. Thermal infrared imaging: detection versus morphometry

The automated pipeline systematically underestimated body length on thermal imagery (bias −19.9 cm, RMSE = 24.1 cm), an offset that was reproducible across surveys and independent of substrate (Section 3.4.1), indicating a consistent methodological cause rather than random noise. Importantly, both the reference (manual) and the automated lengths were measured on the same LWIR orthomosaic, so the bias is not an absolute sensor error relative to a true length: boundary blur common to both measurements would cancel in their difference. The bias instead reflects a systematic difference in how the segmentation model and the human annotator treat low-contrast thermal edges. At the GSD of the M3T at 20 m (∼3.2 cm/px for LWIR versus ∼0.5 cm/px for RGB, consistent with the roughly six-fold RGB–thermal GSD gap reported by Larsen et al. [35]), a ∼120 cm seal spans only ∼38 thermal pixels along its major axis, and thermal dissipation smears the weakly emitting snout and tail into the surrounding background. The segmentation model places the body contour where the thermal gradient collapses and therefore truncates these extremities, whereas the human annotator extrapolates the full outline from anatomical expectation; the difference accumulates as a one-directional shortening of the automated length. Because the M3T records RGB and LWIR simultaneously, the two contributions can be separated. On the three paired acquisitions in which the same colony was reconstructed in both modalities within the same minute, manual lengths measured on the LWIR orthomosaic averaged 3.5 cm longer than manual lengths on the simultaneous RGB orthomosaic (n = 226 and 222 animals respectively; Mann–Whitney p = 0.03), whereas automated lengths averaged 16.9 cm shorter. The annotator’s extrapolation thus accounts for roughly a sixth of the discrepancy and the model’s truncation for the remainder. Instructing annotators to stop where the thermal gradient collapses would improve reproducibility between observers and would remove the smaller component, but it would do so by moving the reference away from the RGB-derived length rather than by improving the automated measurement: the resulting near-zero bias would be an artefact of a redefined reference, not evidence of accuracy. A meaningful accuracy assessment for thermal morphometry therefore needs an external reference — simultaneous RGB imagery of the same individuals, as here, or physically measured animals — rather than a recalibrated annotation protocol. The coarse LWIR resolution is therefore the enabling condition rather than the proximate cause of the bias, operating through the segmentation–annotation discrepancy described above rather than through any loss of length relative to the true animal. The two contributions act in the same direction and cannot be separated with the present data. Body width showed the same directional pattern (thermal bias −2.9 cm versus −0.7 cm for RGB; Section 3.4.1), as expected if a single edge-localization effect governs both axes.

Thermal drone surveys of pinnipeds are widely used for detection and counting, where reduced resolution is not critical. Seymour et al. [36] demonstrated effective automated counting of gray seals from thermal orthomosaics (2–5% discrepancy with manual counts). Larsen et al. [35] used thermal modality to localize *P. vitulina* and *Callorhinus ursinus* on rocky substrate. Christman et al. [37] evaluated thermal surveys for *P. vitulina* on the Aleutian Islands. None of these studies validated morphometric accuracy of thermal measurements - all were limited to detection.

Our results supply this missing quantitative assessment for *P. vitulina* and define a practical limit: at thermal GSD on the order of 3 cm/px, automated body lengths are systematically underestimated by ∼20 cm relative to manual delineation on the same imagery, which precludes reliable size-based classification from thermal data. This does not diminish the value of thermal surveys for detection and counting under reduced visibility, in dense aggregations, or on low-contrast substrates; it means only that body-size metrics should be derived from RGB imagery.

### 4.4. Size-based classification: capabilities, limitations, and prospects

About four in five seals in the RGB subset were assigned to the correct size category (OA = 78.7%; thermal imagery is excluded throughout Section 3.5). Two points of context are important. First, the “true” class is a size category by manual orthomosaic length, not biological age; all classification errors stem from measurement errors (mean deviation for misclassified individuals: 14–18 cm, exceeding RGB body-length RMSE of 12.5 cm). A potential additional error source is inter-observer variability in manual annotation. Although a formal inter-observer agreement assessment was not conducted in this study, published validations for pinnipeds report ICC > 0.99 and mean inter-observer discrepancy of 1–3 cm [8], an order of magnitude below the pipeline body-length RMSE (∼12.5 cm). This suggests that manual annotation error does not materially affect the classification accuracy estimates reported here. Second, away from class boundaries, classification accuracy depends on the width of uncertainty zones, which is governed by orthomosaic geometric accuracy and thus by CET: at CET ≤ 1.53 m, 80.4% of individuals can be reliably classified; at 1.53 < CET ≤ 3.98 m only 28.5%, and above 3.98 m only 20.2%.

Both boundaries involve biological complications beyond measurement error. The pup/juvenile boundary (90 cm) is complicated by post-weaning fasting: mass loss at roughly constant length decouples body length from developmental stage (and broadens within-class L/W variation), so length-based classification is unreliable near this threshold. The juvenile/subadult boundary (130 cm) is complicated by sexual dimorphism: adult female *P. vitulina* (140–161 cm) [23] can be indistinguishable from large juvenile males. The classification is size-based, not age-based in a strict biological sense: growth variability, the spread of birth dates within the pupping season, and population differences in asymptotic length [12] mean that size category is not equivalent to age. For monitoring purposes, separation into size classes remains a standard approach corresponding to functional demographic categories [13,23,38]. Transition from 2D to 3D volumetric features may improve classification: body volume integrates length, width, and profile height and is less sensitive to posture.

A further measurement caveat concerns body width. Width was taken as the maximum dorsal body width of the trunk rather than as an axillary girth: because the flippers were excluded from segmentation, no fore-flipper landmark was available to anchor a standard axillary position, and the location of the maximum width — and, with it, its magnitude — can shift with body posture. We are currently developing a 3D photogrammetric reconstruction of each individual [39,40], which should allow the chest girth to be measured directly and thereby provide a more reliable body-condition metric than a 2D dorsal width.

### 4.5. Automated versus manual size-class composition

Automated classification of all 2,545 pipeline detections (2,298 TP + 247 FP) yielded the following size-class proportions: 39.5% pups, 51.0% juveniles, 9.5% subadults/adults. The like-for-like comparison is between the RGB subsets: the automated pipeline returned 34.8% pups, 53.2% juveniles and 12.0% subadults/adults (n = 1,937 detections), whereas manual measurement of the same RGB orthomosaics returned 25.0% pups, 66.6% juveniles and 8.4% subadults/adults (n = 1,821). Across all sensors the manual composition was 23.1% pups, 67.8% juveniles and 9.1% subadults/adults (n = 2,483). The discrepancy reflects the systematic effect of measurement error on classification near the class boundaries, and it depletes the juvenile class in both directions rather than in one. Among the 1,743 RGB true positives, 147 true juveniles were assigned to pups (mean underestimation −18.2 cm) and 120 to subadults/adults (mean overestimation +15.4 cm), whereas only 40 true subadults/adults were assigned to juveniles (Section 3.5.1; Fig 11). The exchange at the upper boundary is therefore strongly asymmetric, and the automated subadult/adult class is inflated rather than depleted: 229 individuals were assigned to it against 148 in the manual reference, an excess of about 55%. Both outer classes are thus over-populated at the expense of the juvenile class, and the compositional bias is a boundary effect in both directions, not a net downward shift.

**Fig 11.**
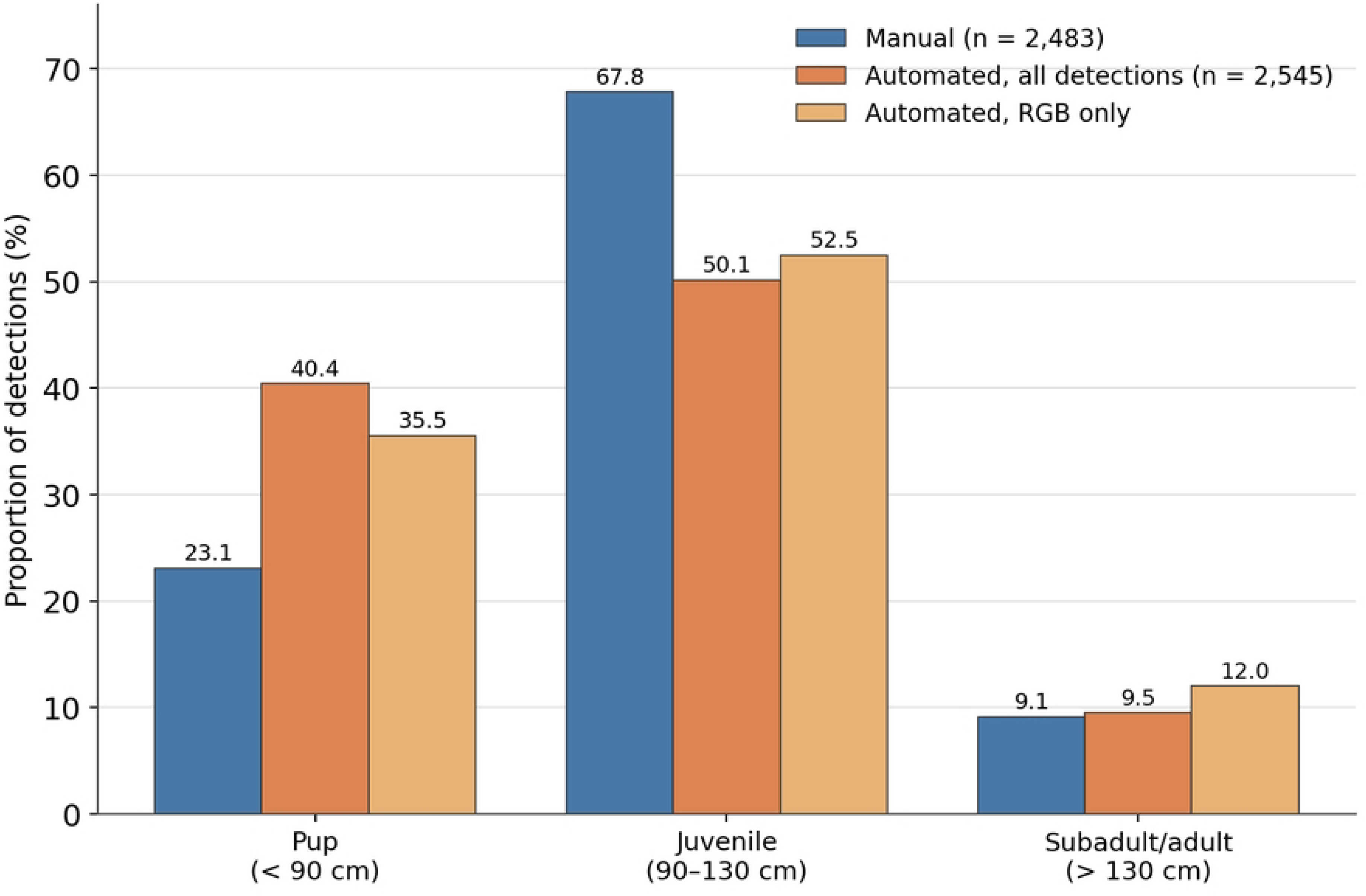
Size-class composition of harbor seals from manual versus automated measurements (RGB; 90 and 130 cm class boundaries). Bars show the percentage of individuals in each size class for manual annotations (all sensors, n = 2,483; RGB only, n = 1,821), all automated pipeline detections (n = 2,545) and automated RGB-only detections (n = 1,937). Automated length error depletes the juvenile class in both directions, inflating the pup class and, to a lesser extent, the subadult/adult class relative to the manual RGB reference.

This composition bias is the population-level manifestation of the boundary misclassification that the classification uncertainty zones characterize at the level of the individual (Section 3.5.2, Table 1). Because these zones—each class boundary ± 2 × the geometric RMSE on the calibration rods—encompass 61.6% of individuals in the aggregate dataset and capture the bulk of misclassified animals, the direction and approximate magnitude of the compositional shift, a net transfer out of the juvenile class across both boundaries that inflates the pup class and, more weakly, the subadult/adult class, are precisely what the zone structure predicts, and the shift grows as orthomosaic geometric quality degrades and the zones widen with increasing CET. The uncertainty zones cannot, however, be used directly as a filter to recover an unbiased composition, because the size classes are themselves concentrated at the boundaries: the pup length range (∼72–90 cm; S2 Table) falls almost entirely within the aggregate 77–103 cm pup/juvenile zone, so excluding zone members would remove most genuine pups and return a non-representative subsample rather than a corrected estimate.

These results confirm that at the current measurement accuracy (RGB body-length RMSE ∼12.5 cm), automated size-class proportions should be interpreted with caution near class boundaries and will benefit from CET-stratified bias correction. A full size-class composition analysis that propagates this classification-uncertainty framework (Section 3.5.2) into the demographic estimates—including CET-stratified bias correction and explicit treatment of the boundary uncertainty zones—together with the detailed demographic interpretation of the observed age composition (seasonal variation, haulout correction factors, and comparison with published population parameters), will be presented in a companion study applying the validated pipeline to the full dataset from the Strait of Georgia (I. A. Usatov and colleagues, manuscript in preparation).

### 4.6. Practical recommendations

Based on our stratified validation results, we propose the following practical recommendations for UAV-based pinniped morphometry and size-class classification.

1. Platform selection. The M3T in RGB mode at 20 m delivered the best overall performance: platform-level orthomosaic geometric accuracy of RMSE = 1.0 cm (the lowest of any platform; Table 2), body-length MAE ≈ 8.6 cm, and detection F1 = 0.967 on slopes. Its factory lens calibration and low (20 m) flight altitude provided scale-invariant geometric accuracy, making it the recommended platform for quantitative morphometry. Lightweight platforms (Mn2, Mn3) are suitable for detection and counting (F1 = 0.91) but yield higher body-length measurement error (MAE 10–11 cm versus 8.6 cm; Table 2).
2. Substrate considerations. Substrate was not a significant predictor of orthomosaic geometric accuracy at the orthomosaic level (Kruskal–Wallis H = 6.27, p = 0.18); the apparent differences among substrates are confounded with platform and flight altitude. On an exploratory basis, flat substrates (plateaus, boulders) yielded somewhat higher classification accuracy than slopes (OA 81.8–82.1% vs. 76.5%), and plateaus also gave the highest detection F1 (0.962 vs. 0.938 on slopes), consistent with distortion of the dorsal projection on inclined surfaces. The pattern is not uniform, however: reefs performed comparably to the flat substrates for classification (OA 80.0%) while returning the lowest detection F1 (0.863), and the one white-sand-beach survey (n = 54 individuals) gave the lowest classification accuracy of any substrate (75.9%), so surface inclination is evidently not the only substrate property that matters. Where surveys must be conducted on slopes, the M3T is therefore preferred for its fine GSD (∼0.5 cm/px at 20 m), and researchers should report substrate type alongside measurement accuracy to enable meaningful cross-study comparison.
3. Classification reliability. Restrict size-based classification to RGB imagery collected at 20 m altitude with platforms providing orthomosaic geometric accuracy (RMSE) < 5 cm on the calibration rods for that survey. Designate the uncertainty bands as each size-class boundary ± 2 × the geometric RMSE measured on the calibration rods; for the aggregate dataset these bands span 77–103 cm (pup/juvenile) and 117–143 cm (juvenile/subadult) and widen for surveys with higher CET (Table 1). We do not recommend assigning individuals within these zones to a single class without additional criteria or post-hoc visual review. Outside these uncertainty zones, classification accuracy reaches 90.9% (cluster-bootstrap 95% CI 87.7–93.7%).
4. Bias correction. Given the systematic nature of platform-specific measurement bias, researchers may apply empirical correction to automated body-length estimates: the M3T tends to overestimate by +1.0 to +3.0 cm, while lightweight platforms (Mn2, Mn3) underestimate by −0.7 to −3.5 cm (RGB mode; these values come from the automated-versus-manual comparison of seal body lengths). These biases are not constant across substrates — for the Mini platforms they run from −0.7 cm on boulders to −3.5 cm on slopes — so a correction should be derived and validated for the specific platform–substrate combination rather than applied at platform level.
5. Thermal imagery should not be used for morphometric measurements of pinnipeds. Thermal data remain valuable for detection and counting, especially in low-visibility conditions or on low-contrast substrates, but the systematic underestimation of automated lengths relative to manual delineation (−19.9 cm, MAE ∼19% of body length) precludes quantitative size-based classification from thermal imagery.
6. For long-term monitoring programs, we recommend deploying calibration rods of known length on haulout substrates prior to the surveys. Calibration rods enable site-specific and survey-specific geometric error assessment, which is essential for detecting temporal trends in body size and age composition - a key objective of ongoing pinniped monitoring in the North Pacific.

## 5. Conclusion

This study presents, to our knowledge, the first pipeline integrating detection, morphometric measurement, and three-class size-based classification of *P. vitulina* from drone orthomosaics in a single automated system. Cascaded DeepLabV3+ segmentation achieved high detection rates under heterogeneous field conditions (four platforms, five substrates, two sensor modalities). Orthomosaic geometric accuracy ranged from 1.0 to 18.0 cm RMSE, with flight altitude (and the integrative CET metric) the significant driver of geometric accuracy, while platform and substrate effects were confounded and did not reach significance at the orthomosaic level. The M3T in RGB mode at 20 m provided the most accurate morphometric performance (geometric RMSE = 1.0 cm; Table 2). Thermal imagery was shown unsuitable for automated pinniped morphometry because automated lengths were systematically shorter than manual delineations on the same imagery. A methodological limitation is that altitude and platform effects were confounded for the M2P (tested only at 40 m); future work should decouple these factors by testing multiple platforms at the same altitude. Threshold classification by body length allowed reliable size-class assignment outside uncertainty zones but yielded moderate accuracy for boundary sizes. The presented pipeline enables a shift from manual counting to automated demographic monitoring with quantitatively characterized uncertainty. Extension to 3D morphometry from the same SfM data is a logical next step and may overcome the key 2D limitation - projection artifacts on sloped substrates.

## Acknowledgments

We thank the internal reviewer at the Marine Mammal Laboratory, Alaska Fisheries Science Center, National Marine Fisheries Service, whose comments on an earlier draft improved the description of the size-class criteria, the presentation of the processing pipeline and the survey-level accuracy results. The present analytical pipeline was developed as a methodological extension of the NPRB-funded pinniped monitoring project (Award NA19NMF4720269) and supported in part by a Marine Mammal Commission grant (MMC25-098). Views and conclusions in this article represent the views of the authors but do not necessarily represent findings or policy of the U.S. National Oceanic and Atmospheric Administration. Any use of trade, firm, or product names is for descriptive purposes only and does not imply endorsement by the U.S. Government.

## Supporting information

**S1 Fig. Water-removal model (n = 800 pairs).** (A) Dice loss; (B) DSC (inset: enlarged scale); (C) learning rate. Best validation DSC = 0.960 at epoch 28; early stopping at epoch 38. LR reduced three times (×0.2).

**S2 Fig. First-level segmentation (n = 4,101 pairs).** (A) Dice loss; (B) DSC; (C) learning rate. Best DSC = 0.911 at epoch 46 of 50. LR reduced three times (×0.2). No train–validation divergence.

**S3 Fig. Second-level segmentation (n = 2,757 pairs).** (A) Dice loss; (B) DSC; (C) learning rate. Best DSC = 0.939 at epoch 48 of 50. LR reduced once (×0.2).

**S1 Table. Calibration and seal-survey dataset: photogrammetric reconstruction-quality indicators and rod measurement-error metrics.** The table has three sections: (1) 22 calibration surveys with control rods, included in the statistical analysis; (2) one calibration survey excluded by the coverage-area criterion (< 500 m²; italic, light-grey shading), because an anomalously high camera density (6,566 cameras ha⁻¹) made its accuracy estimate unrepresentative; and (3) 31 seal surveys with measured animals but no control rods, listed to show the comparability of photogrammetric-condition ranges between the calibration and working datasets. One orthomosaic (Mn3, site H0054, 2023-05-03, 08:20, reefs) carried both calibration rods and measured seals and is therefore listed once in section (1) and once in section (3), with identical reconstruction metrics in both rows. Columns are grouped as survey metadata, reconstruction quality, calibration rods (n, MAE, RMSE), and seal counts. For each survey the number of seals is given both as Manual (number of seals annotated by hand, true positives + false negatives) and Automated (number detected by the pipeline, true positives + false positives). CET (total camera location error, m) is the root-mean-square camera-position residual after block adjustment and is the principal predictor of geometric accuracy (Section 3.3). Bold marks CET > 5 m and rod RMSE > 10 cm, above the typical range. Median rows are given separately for the calibration and seal datasets. MAE, mean absolute error; RMSE, root-mean-square error of rod-length measurement on the orthomosaic (not applicable to seal surveys, dash). UAV codes: M3T, DJI Mavic 3 Thermal; M2P, DJI Mavic 2 Pro; Mn2, DJI Mini 2; Mn3, DJI Mini 3 Pro.

**S2 Table. Summary of body length and body mass of *P.*** *vitulina* by age class (from published data).

**S3 Table. Detection accuracy by platform–sensor–substrate–altitude combination (n ≥ 40).**

**S1 Dataset. Per-animal morphometric measurements.** One row per manually annotated animal and per pipeline detection across the 31 seal-survey orthomosaics (n = 2,720). Columns give the orthomosaic, the detection status of the manual and the automated record, manual and automated body length and width (m), and the platform, flight altitude, sensor and substrate of the survey. All detection, measurement and classification results in Sections 3.2, 3.4, 3.5 and 3.6 are derived from this file.

**S2 Dataset. Calibration-rod measurements.** One row per calibration-rod measurement (n = 601) giving the true and the orthomosaic-measured rod length (m), the orthomosaic, and the platform, flight altitude and substrate of the survey. The geometric-accuracy results in Section 3.3 use the 545 measurements from the 22 retained calibration orthomosaics.

## References

1. Johnston DW. Unoccupied aircraft systems in marine science and conservation. Annu Rev Mar Sci. 2019;11: 439–463. 10.1146/annurev-marine-010318-095323

2. Fiori L, Doshi A, Martinez E, Orams MB, Bollard-Breen B. The use of unmanned aerial systems in marine mammal research. Remote Sens. 2017;9(6): 543. 10.3390/rs9060543

3. Álvarez-González M, Suarez-Bregua P, Pierce GJ, Saavedra C. Unmanned aerial vehicles (UAVs) in marine mammal research: A review of current applications and challenges. Drones. 2023;7(11): 667. 10.3390/drones7110667

4. Larsen GD, Johnston DW. Growth and opportunities for drone surveillance in pinniped research. Mammal Rev. 2024;54(1): 1–12. 10.1111/mam.12325

5. Boveng PL, Bengtson JL, Withrow DE, Cesarone JC, Simpkins MA, Frost KJ, et al. The abundance of harbor seals in the Gulf of Alaska. Mar Mamm Sci. 2003;19(1): 111–127. 10.1111/j.1748-7692.2003.tb01096.x

6. Infantes E, Carroll D, Silva WTAF, Härkönen T, Edwards SV, Harding KC. An automated work-flow for pinniped surveys: A new tool for monitoring population dynamics. Front Ecol Evol. 2022;10: 905309. 10.3389/fevo.2022.905309

7. Dujon AM, Ierodiaconou D, Geeson JJ, Arnould JPY, Allan BM, Katselidis KA, et al. Machine learning to detect marine animals in UAV imagery: effect of morphology, spacing, behaviour and habitat. Remote Sens Ecol Conserv. 2021;7(3): 341–354. 10.1002/rse2.205

8. Hodgson JC, Holman D, Terauds A, Koh LP, Goldsworthy SD. Rapid condition monitoring of an endangered marine vertebrate using precise, non-invasive morphometrics. Biol Conserv. 2020;242: 108402. 10.1016/j.biocon.2019.108402

9. Carroll D, Infantes E, Pagan EV, Harding KC. Approaching a population-level assessment of body size in pinnipeds using drones, an early warning of environmental degradation. Remote Sens Ecol Conserv. 2025;11(2): 156–171. 10.1002/rse2.413

10. Rode KD, Fischbach A, Synnott M, Stewart J, Northcraft N, Allen E, Trotto K, Vancsok C, Issenjou N, Ploof S, Rager S, DiRocco S, Owens S, Prahl A. Photogrammetry-based body condition for monitoring an Arctic marine mammal experiencing habitat loss. Mar Ecol Prog Ser. 2024;751: 211–227. 10.3354/meps14738

11. Trukhanova IS, Altukhov AV, Burkanov VN. Spotted seal (Phoca largha) abundance and size composition at summer haulouts in the southwestern Bering Sea. Aquat Mamm. 2026;52(1): 54–69. 10.1578/AM.52.1.2026.54

12. Harding KC, Salmon M, Teilmann J, Dietz R, Härkönen T. Population wide decline in somatic growth in harbor seals-early signs of density dependence. Front Ecol Evol. 2018;6: 59. 10.3389/fevo.2018.00059

13. Merkel B, Lydersen C, Yoccoz NG, Kovacs KM. The world’s northernmost harbour seal population-How many are there? PLoS ONE. 2013;8(7): e67576. 10.1371/journal.pone.0067576

14. Majewski SP, Ellis GM. Abundance and distribution of Harbour Seals (Phoca vitulina) in the Strait of Georgia, British Columbia. DFO Can Sci Advis Sec Res Doc. 2022/060. 2022.

15. Tucker S, Majewski S, Nordstrom C, Trzcinski MK, Doniol-Valcroze T. Stock assessment of Pacific harbour seals (Phoca vitulina richardii) in Canada in 2019. DFO Can Sci Advis Sec Res Doc. 2025/069. v + 82 p. 2025. https://open-science.canada.ca/handle/123456789/4062

16. Olesiuk PF. An assessment of the status of harbour seals in British Columbia. DFO CSAS Res Doc. 1999/033. 1999.

17. Teilmann J, Rigét FF, Härkönen T. Optimizing survey design for Scandinavian harbour seals. ICES J Mar Sci. 2010;67(5): 952–958. 10.1093/icesjms/fsq001

18. Härkönen T, Heide-Jørgensen M-P. Age- and sex-specific behaviour in harbour seals leads to biased estimates of vital population parameters. J Appl Ecol. 1999;36(1): 36–45. 10.1046/j.1365-2664.1999.00434.x

19. Hamilton CD, Lydersen C, Ims RA, Kovacs KM. Haul-out behaviour of harbour seals (Phoca vitulina) throughout the year. PLoS ONE. 2014;9(1): e86055. 10.1371/journal.pone.0086055

20. Cottrell PE, Jeffries S, Beck B, Ross PS. Growth and development in free-ranging harbor seal (Phoca vitulina) pups from southern British Columbia, Canada. Mar Mamm Sci. 2002;18(3): 721–733. 10.1111/j.1748-7692.2002.tb01069.x

21. Rapado-Tamarit B, Méndez-Aróstegui M, de Reus K, Sarraude T, Pen I, Groothuis TGG. Age estimation and growth patterns in young harbor seals (Phoca vitulina vitulina) during rehabilitation. J Mammal. 2025;106(2): 491–504. 10.1093/jmammal/gyae128

22. Hauksson E. Growth and reproduction in the Icelandic common seal. Mar Biol Res. 2006;2(1): 59–73. 10.1080/17451000600650038

23. Lydersen C, Kovacs KM. Growth and population parameters of harbour seals in Svalbard, Norway. Polar Biol. 2005;28(2): 156–163. 10.1007/s00300-004-0656-7

24. Bierlich KC, Schick RS, Hewitt J, Dale J, Goldbogen JA, Friedlaender AS, et al. Bayesian approach for predicting photogrammetric uncertainty in morphometric measurements derived from drones. Mar Ecol Prog Ser. 2021;673: 193–210. 10.3354/meps13814

25. Currie JJ, Stirling B, Barber-Meyer S. Validation of the DJI Mavic 3 Pro for field-based whale photogrammetry. Aquat Mamm. 2026;52(2): 175–182. 10.1578/AM.52.2.2026.175

26. Dawson SM, Bowman MH, Leunissen E, Sirguey P. Inexpensive aerial photogrammetry for studies of whales and large marine animals. Front Mar Sci. 2017;4: 366. 10.3389/fmars.2017.00366

27. American Society of Mammalogists. Mammalian measurements. J Mammal. 1967;48(4): 733–734.

28. Chen L-C, Zhu Y, Papandreou G, Schroff F, Adam H. Encoder-decoder with atrous separable convolution for semantic image segmentation. In: ECCV. 2018. pp. 801–818. 10.1007/978-3-030-01234-2_49

29. Dice LR. Measures of the amount of ecologic association between species. Ecology.1945;26(3): 297–302. 10.2307/1932409

30. Milletari F, Navab N, Ahmadi S-A. V-Net: Fully convolutional neural networks for volumetric medical image segmentation. In: 2016 Fourth International Conference on 3D Vision (3DV). IEEE; 2016. pp. 565–571. 10.1109/3DV.2016.79

31. Sudre CH, Li W, Vercauteren T, Ourselin S, Cardoso MJ. Generalised Dice overlap as a deep learning loss function for highly unbalanced segmentations. In: DLMIA/ML-CDS 2017. Springer; 2017. pp. 240–248. 10.1007/978-3-319-67558-9_28

32. Goodfellow I, Bengio Y, Courville A. Deep learning. Cambridge, MA: MIT Press; 2016.

33. Pérez Tadeo M, Gammell M, O’Callaghan SA, O’Connor I, O’Brien J. Disturbances due to unmanned aerial vehicles (UAVs) on harbor seal (Phoca vitulina) colonies: recommendations on best practices. Mar Mamm Sci. 2023;39(3): 757–779. 10.1111/mms.13002

34. Liu J, Jin F, Ji M, Qu L, Wang J, Wang C. Hierarchical dual-model detection framework for spotted seals using deep learning on UAVs. Animals. 2025;15(21): 3100. 10.3390/ani15213100

35. Larsen GD, Seymour AC, Richmond EL, Divine LM, Moreland EE, Newton E, et al. Drones reveal spatial patterning of sympatric Alaskan pinniped species and drivers of their local distributions. Drone Syst Appl. 2022;10(1): 235–255. 10.1139/dsa-2021-0050

36. Seymour AC, Dale J, Hammill M, Halpin PN, Johnston DW. Automated detection and enumeration of marine wildlife using unmanned aircraft systems (UAS) and thermal imagery. Sci Rep. 2017;7: 45127. 10.1038/srep45127

37. Christman CL, London JM, Conn PB, Hardy SK, Brady GM, Dahle SP, et al. Evaluating the use of thermal imagery to count harbor seals in aerial surveys. Mamm Biol. 2022;102: 719–732. 10.1007/s42991-021-00191-6

38. Boulva J, McLaren IA. Biology of the harbor seal, Phoca vitulina, in eastern Canada. Bull Fish Res Board Can. 1979;200: 1–24.

39. Usatov IA, Trukhanova IS, Burkanov VN, Altukhov AV, Bowker A, Nordstrom C, et al. Preliminary results on 3D morphometry of pinnipeds to improve body condition assessment and counts from UAV imagery data. Presented at: Alaska Marine Science Symposium; 2026 Jan; Anchorage, AK. 2026.

40. Usatov IA, Trukhanova IS, Burkanov VN, Altukhov AV, Bowker A, Nordstrom C, et al. Factors affecting the success of 3D reconstruction of pinnipeds from UAV data: effects of platform type, flight altitude, and imaging mode. Accepted for: 26th Biennial Conference on the Biology of Marine Mammals; 2026; San Juan, Puerto Rico. 2026.

